# Uncovering transcriptional regulatory network during regeneration for boosting wheat transformation

**DOI:** 10.1101/2022.10.21.513305

**Authors:** Xuemei Liu, Xiaomin Bie, Xuelei Lin, Menglu Li, Hongzhe Wang, Xiaoyu Zhang, Yiman Yang, Chunyan Zhang, Xiansheng Zhang, Jun Xiao

## Abstract

Genetic transformation is important for gene functional study and crop breeding. Though it is available in many plant species, the transformation efficiency in wheat is generally low, which greatly restricts the genetic manipulation in wheat. Here, we use multi-omic analysis strategy to uncover core transcriptional regulatory network (TRN) driving wheat shoot regeneration and identify key factors that boost the transformation efficiency. RNA-seq, ATAC-seq and CUT&Tag were used to profile the transcriptome and chromatin dynamic during regeneration process from immature embryo of wheat variety Fielder. Sequential expression of gene clusters that mediating cell fate transition during regeneration is induced by auxin signaling, in coordination with changes of chromatin accessibility, H3K27me3 and H3K4me3 status. The TRN driving wheat shoot regeneration was built-up and 446 key transcriptional factors (TFs) occupied the core of network were identified, including functionally tested regeneration factors in other species. We further compared the regeneration process between wheat and *Arabidopsis* and found that DNA binding with one finger (DOF) TFs show distinct patterns in two species. Furthermore, we found that *TaDOF5*.*6* (*TraesCS6A02G274000*) and *TaDOF3*.*4* (*TraesCS2B02G592600*) can significantly improve the transformation efficiency of different wheat varieties. Thus, our data uncovers the molecular regulatory insights for wheat shoot regeneration process and provides potential novel targets for improving transformation efficiency in wheat.

## Introduction

Since Gottleib Haberland proposed totipotency of plant cells in 1902 (Haberlandt, 2003), plant tissue culture technology has been widely used in various fields, especially in the acquisition of transgenic and gene-edited plants (Bidabadi and Jain, 2020; Thorpe, 2007). Conventional shoot regeneration includes two stages: callus induction on auxin rich callus-inducing medium (CIM) and subsequent shoot induction on cytokinin rich shoot-inducing medium (SIM), which is also called *de novo* shoot organogenesis (DNSO) (Radhakrishnan et al., 2018; Shin et al., 2019). Almost all organs such as leaves, hypocotyls and protoplasts of the eudicot *Arabidopsis thaliana* can be induced to regenerate, while as in monocot cereals for instance wheat, only certain undifferentiated tissues could be induced to form callus, such as immature embryos (Hayta et al., 2019; Ikeuchi et al., 2019). This maybe because the mature tissues of cereals no longer retain regeneration-competent cells that retained in mature organs of *Arabidopsis* (Hu et al., 2017).

Studies in *Arabidopsis* suggest that the acquisition of totipotency on CIM follows a lateral root initiation pathway, regardless of the origin of explants (Atta et al., 2009; Sugimoto et al., 2010). At the molecular level, auxin signaling induces *WUSCHEL RELATED HOMEOBOX 11* (*WOX11*) and *WUSCHEL RELATED HOMEOBOX 12* (*WOX12*) expression and transform regeneration-competent cells into founder cells, which then activates *WUSCHEL RELATED HOMEOBOX 5* (*WOX5*) and *LATERAL ORGAN BOUNDARIES DOMAIN 16* (*LBD16*) to promote the transformation of founder cells into root primordia (Hu and Xu, 2016; Liu et al., 2018b, 2014). When callus are transferred on SIM, the root stem cell genes *PLETHORA 1* (*PLT1*), *PLETHORA 2* (*PLT2*), *SCARECROW* (*SCR*) and *ROOT-CLAVATA HOMOLOG1* (*RCH1*) are rapidly down regulated. While genes related to shoot meristem, such as *CUP SHAPED COTYLEDON1* (*CUC1*), *CUP SHAPED COTYLEDON2* (*CUC2*), *SHOOT MERISTEMLESS* (*STM*) and *WUSCHEL* (*WUS*), are activated to promote callus transform into shoot meristem (Atta et al., 2009; Gordon et al., 2007; Meng et al., 2017). *PLETHORA 3* (*PLT3*), *PLETHORA 5* (*PLT5*) and *PLETHORA 7* (*PLT7*) of AP2 family not only participate in callus formation by regulating root stem cell regulators *PLT1* and *PLT2*, but also regulate *CUC1* and *CUC2* on SIM to mark the pre-meristematic region (Kareem et al., 2015).

Besides key TFs, some epigenetic regulators have also been identified that are involved in DNSO through crosstalk with auxin or cytokinin signals in *Arabidopsis*. The reprogramming of H3K27me3 is critical for callus formation, especially the loss of H3K27me3 at auxin pathway genes and root regulatory genes (He et al., 2012). H3K36me3 methyltransferase *ARABIDOPSIS TRITHORAX-RELATED 2* (*ATXR2*) interacts with *AUXIN RESPONSE FACTOR 7* (*ARF7)* and *ARF19* to promote the expression of *LBD16* and *LBD29* during callus induction (Lee et al., 2017), while on SIM, *ATXR2* interacts with type-B *ARABIDOPSIS RESPONSE REGULATOR 1* (*ARR1*) to activate type-A ARR, thereby inhibiting the expression of *WUS* and affecting shoot regeneration (Lee et al., 2021). In contrast, the methyltransferase of H3K4me3 *ARABIDOPSIS TRITHORAX 4* (*ATX4*) inhibits callus induction but promote shoot regeneration (Lee et al., 2019). In addition, DNA methyltransferase DNA *METHYLTRANSFERASE1* (*MET1*) is regulated by cytokinin-CYCD3-E2FA module and inhibits the expression of *WUS* in callus (Liu et al., 2018a).

Compared to *Arabidopsis*, cereals are recalcitrant to regeneration (Ikeuchi et al., 2019). In wheat, immature embryo is widely used as explant for regeneration, which limits the application of transformation (Hayta et al., 2019; Hiei et al., 2014). Besides, different wheat genotypes show varied regeneration efficiency, in particular some agronomic important varieties have low regeneration efficiency (Zhang et al., 2018; Wang et al., 2017). Recently, some ‘regeneration factors’ identified in *Arabidopsis* have been applied into cereals to improve the efficiency of genetic transformation. Overexpression of *BABY BOOM* (*BBM*) and *WUSCHEL 2* (*WUS2*) can promote the transformation efficiency of a variety of cereals including maize, rice and sorghum (Lowe et al., 2016; Suo et al., 2021). *WOX5* significantly increased the transformation efficiency of multiple wheat varieties in a genotype-independent manner (Wang et al., 2022). The fusion proteins of *GROWTH REGULATING FACTOR 4*-*GROWTH REGULATING FACTOR INTERACTING FACTOR 1*(*GRF4-GIF1*) can also enhance the transformation efficiency and improve genome editing of wheat (Debernardi et al., 2020; Qiu et al., 2022). However, the molecular mechanism underlies regeneration process in cereals is not clear. Considering the regeneration capacity difference between monocot and edict, one would speculate that conserved and species-specific key factors might be involved.

Here, taking hexaploid wheat as an example, we captured transcription and chromatin dynamics during shoot regeneration process. By comparison with *Arabidopsis*, similar and wheat-specific transcriptional and epigenetic features were identified. Based on the high correlation of chromatin accessibility and gene expression, we established a TRN for driving wheat shoot regeneration. A total of 446 core TFs were identified, among which *TaDOF5*.*6* and *TaDOF3*.*4* significantly improved transformation efficiency of wheat in different varieties.

## Results

### A transcriptional and epigenomic atlas during wheat shoot regeneration

As genetic transformation in wheat is genotype-dependent, we selected the most easily transformed variety Fielder as material for studying the regeneration process (Hayta et al., 2019; Wang et al., 2017). In the process of wheat shoot regeneration, the scutellum from immature embryo of approximately 14 days after anthesis was used as explant. After 3 days of induction, scutellum lost its character and started to cell division and form callus. After 6 days of induction, the callus surface began to protrude and these protrusions became dominant and dense after 9 days, initiating the formation of promeristem of shoot apical meristem (SAM). After 12 days of induction, several layers of cells were exfoliated from the callus and protrusions near callus surface continued to develop, with more SAMs formed (Fig. 1a). Thus, to capture the transcription and chromatin dynamic during wheat shoot regeneration and identify the core regulators, we generated fifteen RNA-seq libraries, eight ATAC-seq libraries and twenty-four CUT&Tag libraries for histone modification (H3K27me3, H3K4me3 and H3K27ac) from five time points during wheat shoot regeneration at 0, 3, 6, 9, 12 days after induction (DAI) (Fig. 1a).

**Fig. 1.**
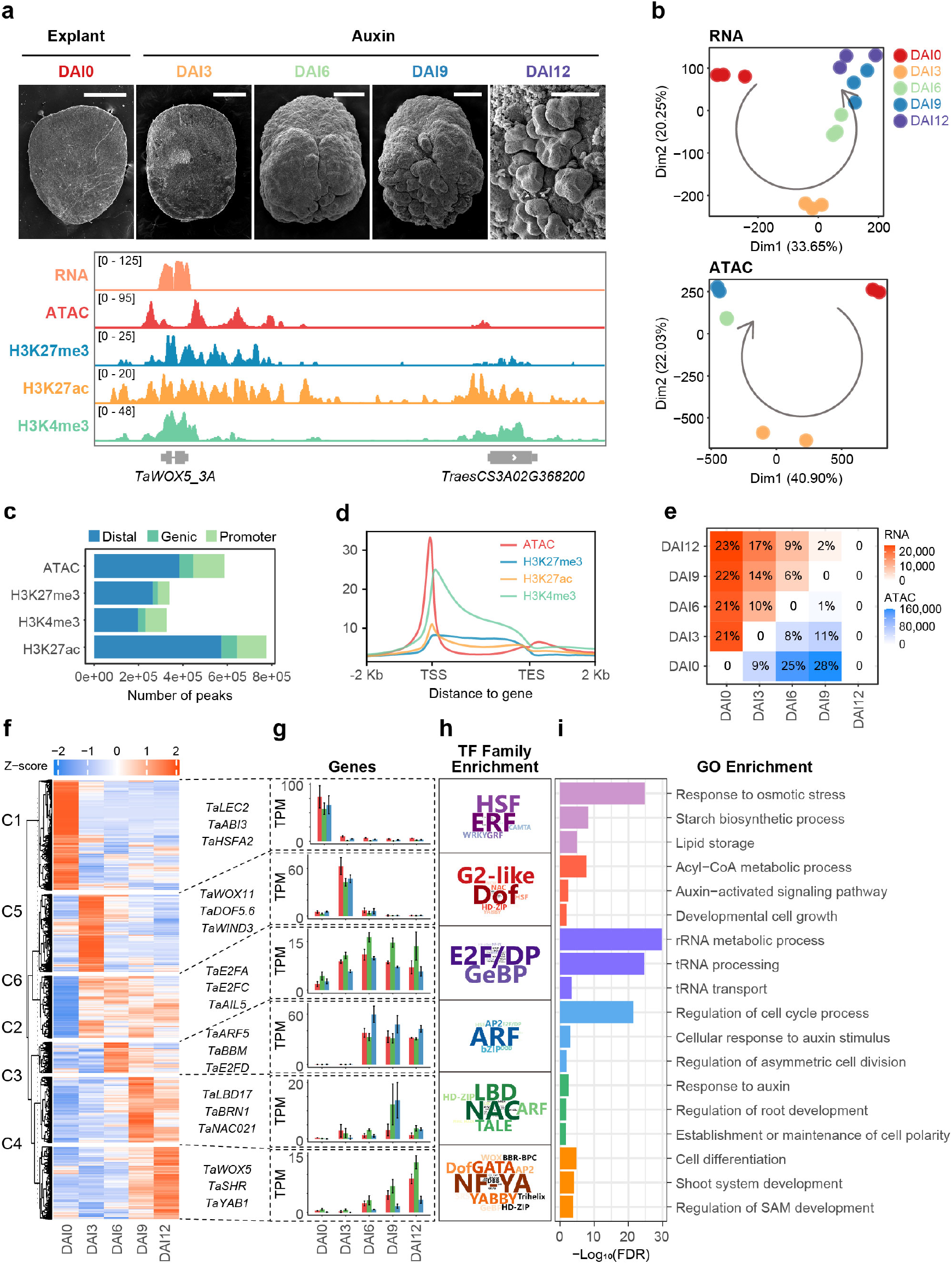
A transcriptional and epigenomic atlas of wheat shoot regeneration. **a**, Morphology of different time points during wheat shoot regeneration for sample collection observed by scanning electron microscopy and screenshots for various data types shown by IGV browser; DAI: day after induction. **b**, Principal component analysis of RNA-seq and ATAC-seq dataset. **c**, Genome-wide distribution of various epigenetic modification peaks. **d**, Profile of epigenetic marks along genic region (data form DAI6 stage); TSS: transcription start site, TES: transcription end site. **e**, Number of differentially expressed genes and differentially accessible peaks among different induction stages; Percentage represents the proportion of differentially expressed genes to all genes or the proportion of differentially accessible peaks to all peaks. **f**, K-means clustering analysis of differentially expressed genes. **g**, Expression level of representative marker genes from each RNA cluster. **h**, TFs family enrichment analysis for each RNA cluster (Fisher’s exact test, p.adj < 1e-6); The front size reflects the degree of enrichment, as indicated by – Log10(p.adj). **i**, Gene ontology enrichment analysis for each RNA cluster.

In general, our transcriptome and epigenome data show high reproducibility between different biological replicates and can reflect the continuity of wheat shoot regeneration process (Fig. 1b, Extended Data Fig. 1a, b). Active chromatin features such as open chromatin, H3K27ac and H3K4me3 are positively correlated with each other, while negatively associated with repressive histone mark H3K27me3 (Extended Data Fig. 1b). Most of the open chromatin and histone modification peaks are located in the intergenic region (Fig. 1c). The distribution profile of open chromatin and three histone modification marks along gene show transcriptional regulation correlated specific patterns (Extended Data Fig. 1c), as previous report (Zhao et al., 2022). The peaks of open chromatin and H3K27ac are mainly near the transcription start site (TSS), while H3K27me3 and H3K4me3 are distributed across the whole gene (Fig. 1d). As well, highly expressed genes are generally in an accessible chromatin state, with more H3K27ac, H3K4me3 and less H3K27me3 (Extended Data Fig. 1c). Conversely, low or no expressed genes marked by H3K27me3 have less active chromatin features (Extended Data Fig. 1c). During the regeneration process, we found 40,146 (37% coding genes of whole genome) differentially expressed genes (DEGs) (Fig. 1e), suggesting an extensive transcriptome reprogramming event. We further identified differentially accessible peaks for ATAC-seq and differentially marked peaks by histone modification. About 38% ATAC-seq peaks are differentially accessible (Fig. 1e), while only 6%, 4% and 1% peaks are differentially marked by H3K4me3, H3K27me3 and H3K27ac, respectively (Extended Data Fig. 1d). This highlights the chromatin accessibility dynamic in shaping wheat shoot regeneration process.

To understand the transcription dynamic during wheat shoot regeneration, we performed K-means clustering analysis and divided DEGs into six clusters (Fig. 1f, Table S1). For each cluster (C1 to C6), transcription factor family enrichment (Fig. 1h, Table S2) and gene ontology enrichment (Fig. 1i, Table S3) were analyzed to determine the underlying biological process. Embryo identity genes such as *LEAFY COTYLEDON 2* (*TaLEC2*) and *ABA INSENSITIVE 3* (*TaABI3*) in C1 are inhibited, which indicates that dedifferentiation of embryo identity is the first requirement for regeneration (Fig. 1g). After auxin treatment, *TaWOX11, WOUND INDUCED DEDIFFERENTIATION 3* (*TaWIND3*) and members in DOF family in C5 show quick and transient activation. By contrast, C6 genes are continuously active upon auxin addition. Genes related to tRNA processing, rRNA metabolic process and cell division are enriched (Fig. 1h, i), indicating cell proliferation and translation is constantly active during the regeneration process. Genes in C2, C3 and C4 are sequentially activated after auxin treatment. C2 contains cell cycle, cell division and auxin response related genes, such as *E2F TRANSCRIPTION FACTOR D* (*TaE2FD*), *BBM* and *AUXIN RESPONSE FACTOR 5* (*TaARF5*) (Fig. 1g). Genes in C3 and C4 are highly expressed in the late stage of wheat callus induction, which are related to root and shoot development, including *TaLBD17, BEARSKIN 1* (*TaBRN1*), NAC *DOMAIN CONTAINING PROTEIN 21*(*TaNAC021*), *WOX5, SHORT ROOT* (*TaSHR*) and *YABBY1* (*TaYAB1*) (Fig. 1g).

Thus, we generated a comprehensive transcriptome and epigenome dataset for wheat shoot regeneration. Both transcription and chromatin accessibility have remarkable changes during this process. In addition, our data also captured the dynamic and sequentially transcriptional alterations that mediating the cell fate transition during this process.

### Chromatin landscape dynamic coordinated with transcriptional profile during wheat shoot regeneration

Gene expression is co-regulated by cis-motif and trans-factors in the context of local chromatin landscape, especially at the regulatory region (Klemm et al., 2019). To explore the correlation between transcriptional profile and chromatin accessibility as well as histone modifications during wheat shoot regeneration, we used two independent approaches. Firstly, we directly calculated the Pearson correlation between transcriptional pattern and various chromatin features for each RNA cluster (Fig. 2a). We found that chromatin accessibility dynamic is highly positive correlated with transcriptional changes in all clusters, while H3K27me3 shows negative correlation in most clusters, especially for C2, C3, C4, C6, with reduction of H3K27me3 accompanied by transcriptional activation (Fig. 2a). For H3K4me3, only C3 and C4, which are highly expressed in late callus induction, show a positive correlation, whereas H3K27ac show a low correlation with transcription in most clusters (Fig. 2a). Representative genes from different clusters are shown to state the correlation between transcription profile and chromatin landscape (Fig. 2b). *TaLEC2* is highly expressed at DAI0, when the chromatin near its gene is accessible, enriched with active H3K27ac and H3K4me3, but not inhibitory H3K27me3. During dedifferentiation, the chromatin of *TaLEC2* is rapidly closed, and the active histone modifications, especially H3K4me3, are reduced, leading to a rapid decline in its expression level. For *TaBBM* and *TaWOX5* activated during callus formation, their high expression in DAI6-9-12 is closely related to the increased chromatin accessibility, the removal of H3K27me3 and the increase of H3K4me3 on them.

**Fig. 2.**
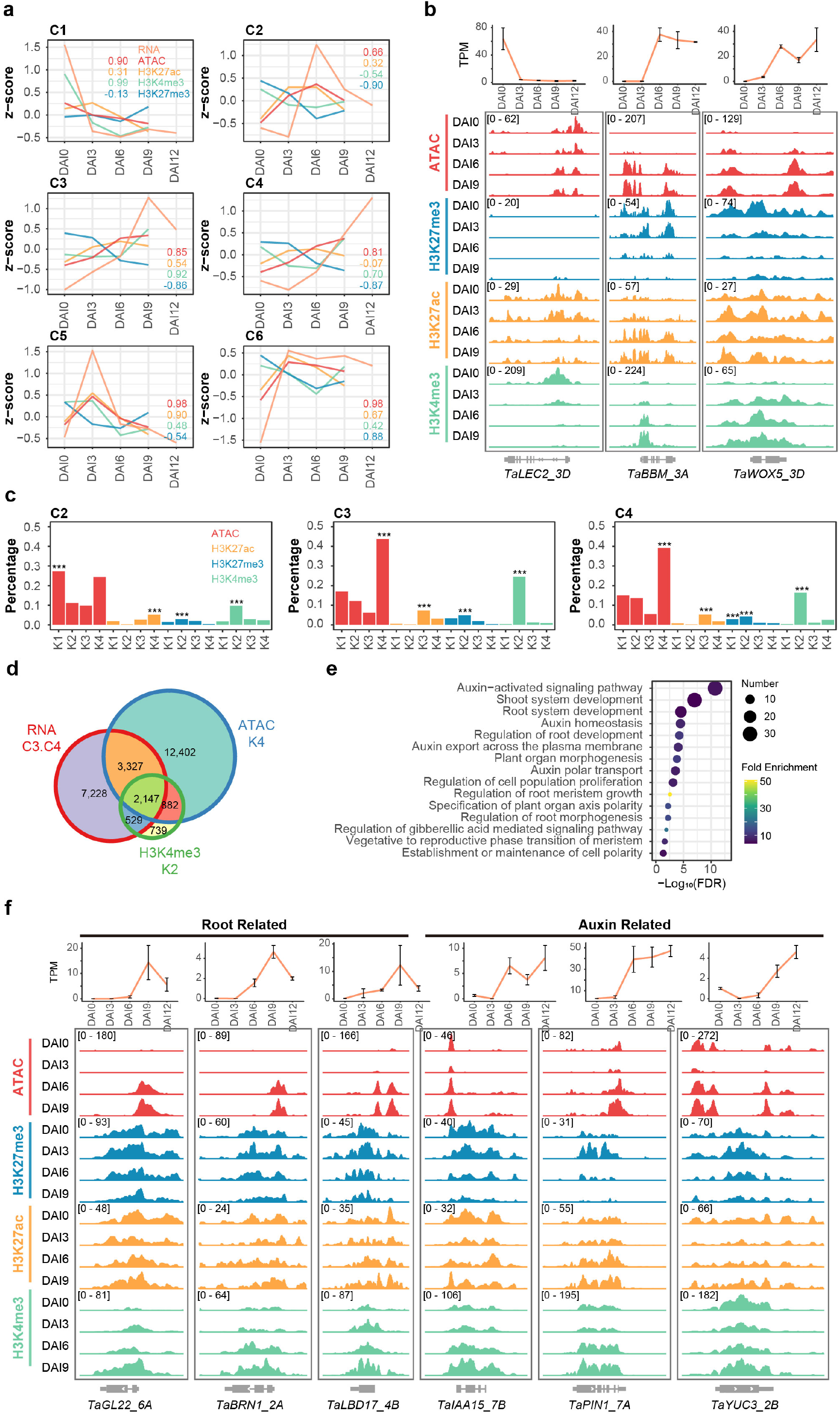
Chromatin dynamic coordinated transcription profile during wheat shoot regeneration. **a**, Correlation analysis between transcription and various epigenetic marks (Pearson correlation analysis). Numbers in different colors indicate the correlation value of individual comparison. **b**, The dynamic transcription and epigenetic modification track for *TaLEC2_3D, TaBBM_3A* and *TaWOX5_3D* during regeneration. **c**, Over-representation analysis for gens in RNA cluster C2, C3 and C4 in ATAC and histone modification cluster (Fisher’s exact test, ***: p<0.001); The percentage represents the proportion of genes that overlap with the epigenetic cluster in the RNA cluster. **d**, Venn diagram for RNA cluster C3, C4, ATAC cluster K4 and H3K4me3 cluster K2. **e**, The gene ontology enrichment analysis of 2,147 intersection genes in d. **f**, The dynamic transcription and epigenetic modification tracks for root related genes *TaGL22_6A, TaBRN1_2A, TaLBD17_4B* and auxin related genes *TaIAA15_7B, TaPIN1_7A, TaYUC3_2B*.

Secondly, we performed K-means clustering analysis for differentially accessible peaks and histone modification marked peaks located in proximal of genes (Extended Data Fig. 2a, Table S4). For each RNA cluster, we searched for the over representation cluster of open chromatin and histone modifications (Table S5). The chromatin accessibility cluster has the most overlap with RNA cluster, followed by H3K4me3, and the least overlap is H3K27me3 and H3K27ac (Fig. 2c, Extended Data Fig. 2b). This suggests that expression of most genes may be regulated by chromatin accessibility, followed by H3K4me3, while H3K27me3 and H3K27ac may coordinate with few genes, which is consistent with the degree of changes in different epigenome data during regeneration (Fig. 1e, Extended Data Fig. 1d). Genes in C3 and C4 activated at late callus induction are significantly enriched in K4 of chromatin accessibility and K2 of H3K4me3 (Fig. 2c). So we took the intersection of them and obtained 2147 genes (Fig. 2d). These genes are involved in root development (Fig. 2e, Table S6), such as *GERMIN-LIKE PROTEIN SUBFAMILY 2 MEMBER 2 PRECURSOR* (*TaGL22*), *TaBRN1, TaLBD17* and auxin signaling pathway (Fig. 2e, Table S6), such as *INDOLE-3-ACETIC ACID INDUCIBLE 15* (*TaIAA15*), *PIN-FORMED 1* (*TaPIN1*) and *YUCCA3* (*TaYUC3*) (Fig. 2e, f). For genes in C2, C3, C4 and C6 with expression changes negatively correlated with H3K27me3 dynamics, are significantly enriched in K1, K2 of chromatin accessibility and K1, K2, K3 of H3K27me3 (Fig. 2c, Extended Data Fig. 2b). A total of 1197 genes (Extended Data Fig. 2c) are regulated by both chromatin accessibility and H3K27me3, including genes involved in auxin signaling pathway (Extended Data Fig. 2d, Table S7), such as *TaIAA15, TaPIN1* and *TaYUC3* (Fig. 2f). Therefore, auxin signaling pathway genes are regulated by chromatin accessibility, H3K4me3 and H3K27me3 simultaneously during wheat shoot regeneration. While root development related genes are mainly regulated by chromatin accessibility and H3K4me3.

In summary, transcription dynamic during wheat shoot regeneration are mainly coordinated with changes in chromatin accessibility and partially with gain of H3K4me3 or loss of H3K27me3 for genes activated during wheat shoot regeneration.

### Auxin signaling promotes callus induction and prepares for the later differentiation via a crosstalk with cytokinin

Hormones play crucial role in plant regeneration, especially auxin and cytokinin. To explore the role of auxin and cytokinin during wheat shoot regeneration, we first divided wheat callus induction into two stages according to the expression pattern of marker genes *TaWOX11* and *TaWOX5* (Fig. 3a). *WOX11* is the identity gene of root founder cell (Liu et al., 2014), and its high expression at DAI3 indicates that regenerative competent cells in wheat embryo have been transformed into root founder cells under the action of auxin at DAI3. At DAI6-9-12, the high expression of *WOX5* (Fig. 3a), a marker gene of root stem cell niche (Sarkar et al., 2007), indicates that the root founder cells have been transformed into root primordium. The formation of the root founder cell requires local auxin maximum. Correspondingly, auxin transport genes, such as *ARABIDOPSIS THALIANA P GLYCOPROTEIN 1* (*TaABCB1*), *ARABIDOPSIS THALIANA P GLYCOPROTEIN 3* (*TaABCB3*) and *LIKE AUXIN RESISTANT 1* (*TaLAX1*) are highly expressed at DAI3 (Fig. 3a), which may be involved in re-distribution of the endogenous auxin in callus. Auxin signal is needed to stimulate the formation and maintain of root primordium (Della Rovere et al., 2013). Accordingly, we found that genes involved in auxin signaling pathway are activated during DAI6-9-12, such as *TRANSPORT INHIBITOR RESPONSE 1* (*TaTIR1*), *TaIAA3* and *TaARF5* (Fig. 3a).

**Fig. 3.**
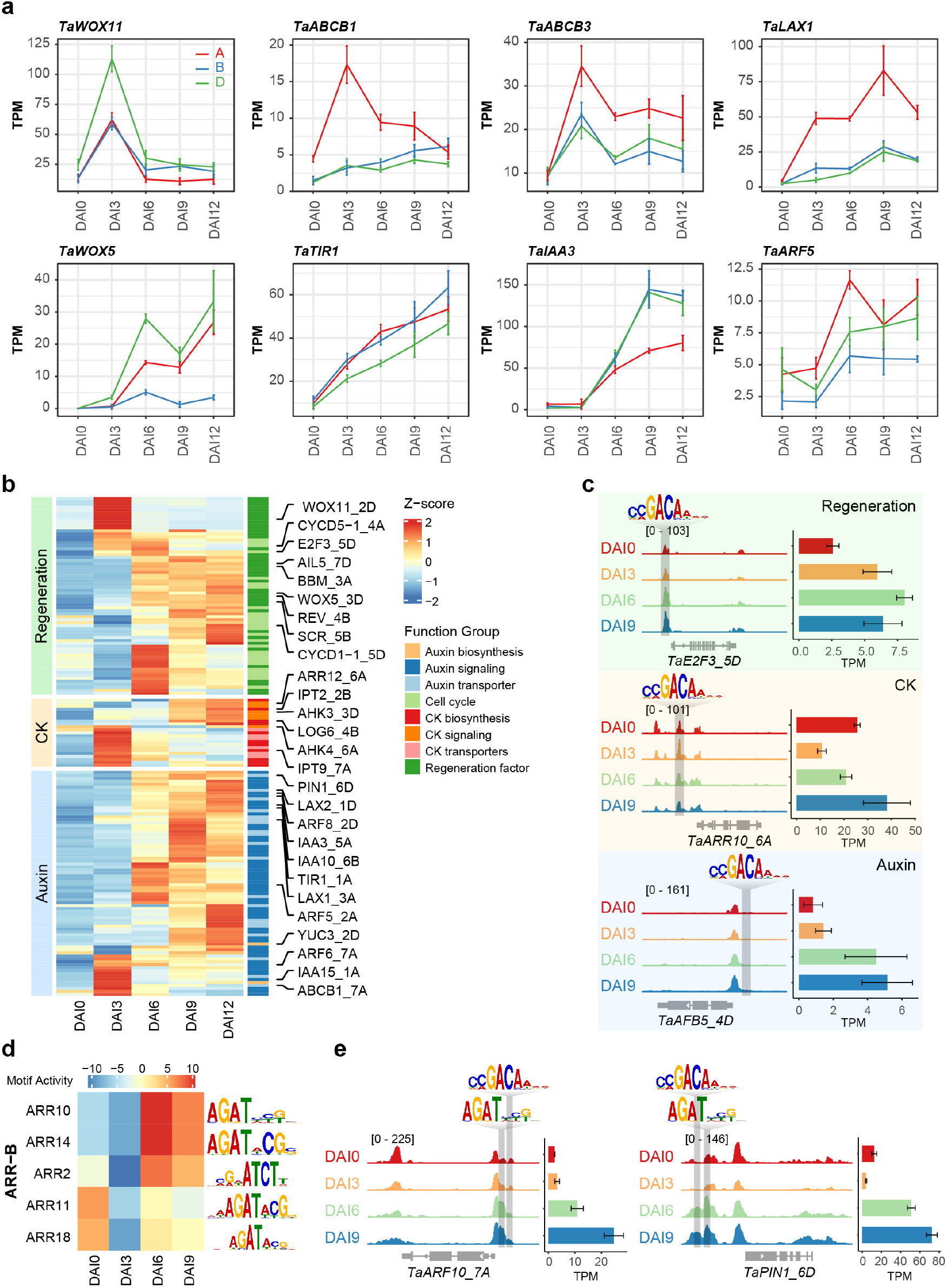
Auxin signaling promoted wheat shoot regeneration. **a**, Expression pattern of *TaWOX11, TaWOX5* and auxin transport and signaling related genes during wheat shoot regeneration. **b**, Expression heatmap for three types of ARFs potential target genes during wheat shoot regeneration. **c**, Presence of AuxRE in the open chromatin region of *TaE2F3_5D, TaARR10_6A* and *TaAFB5_4D*, with dynamically activated expression during regeneration. **d**, Motif activity dynamic for type-B ARR during wheat shoot regeneration. Motif activity represents the chromatin accessibility at the transcription factor binding sites. **e**, Presence of ARFs and type-B ARRs binding motif in the open chromatin region of *TaARF10_7A* and *TaPIN1_6D*, with gradually elevated expression after DAI6.

Next, we wonder how auxin signaling activates regeneration process. ARFs can bind to auxin response elements (AuxRE) and transmit auxin signals to downstream pathways (Weijers and Wagner, 2016). Thus, we identified ARF targets during regeneration process via looking for genes with AuxRE in the proximal open chromatin regions and elevated expression after auxin application (Extended Data Fig. 3a). In total, 9,471 target genes related to auxin signaling pathway, cell cycle and shoot development were identified (Extended Data Fig. 3b, Table S8). Within these genes, three major groups were highlighted, including meristematic identity genes, cytokinin synthesis and signaling genes, and auxin synthesis, transport and signaling pathway genes (Fig. 3b, Table S9). Of note, the chromatin accessibility near AuxRE of these three category genes is relatively higher than random regions and the accessibility is increased with callus induction, while less abundant of H3K27me3 compared to random regions (Extended Data Fig. 3c). The first group includes many known genes involved in plant regeneration, such as meristem identity genes *TaWOX5, TaBBM, TaSCR* and some cell cycle related genes, such as *CYCLIN D1;1* (*TaCYCD1-1)* and *E2F TRANSCRIPTION FACTOR 3* (*TaE2F3)* (Fig. 3c), indicating auxin signaling functions in driving the callus induction process. Genes in the second group are related to cytokinins, including cytokinin synthesis gene *TRNAISOPENTENYLTRANSFERASE 2* (*TaIPT2)*, cytokinin signaling pathway gene *ARABIDOPSIS HISTIDINE KINASE 3* (*TaAHK3)* and *ARABIDOPSIS RESPONSE REGULATOR 10* (*TaARR10)* (Fig. 3c). This highlights that auxin signaling could activate cytokinin pathway for priming the later organ differentiation program. Genes in the third group include auxin synthesis gene *TaYUC3*, transport gene *PIN-FORMED 1* (*TaPIN1*) and signaling pathway gene *AUXIN F-BOX PROTEIN 5* (*TaAFB5*) (Fig. 3c), with elevated expression at different stages during regeneration. Thus, ARFs may play a role in maintaining the balance of auxin concentration and signaling at different stages during wheat shoot regeneration.

It is generally believed that cytokinins determine shoot fate on SIM, whereas recent studies have found that cytokinins, especially B-ARR, also play a role on CIM (Wu et al., 2022). Here, we found that cytokinin related genes may be directly activated by ARFs during callus induction in wheat (Fig. 3b, c), and the motif activity of B-ARR is significantly increased at DAI6, as evaluated by ChromVar (Fig. 3d). Indeed, we found 2,924 potential targets of B-ARR TFs, based on the presence of ARR binding motifs (AGAT) in the open chromatin region of genes with active expression at DAI6 and DAI9 (Extended Data Fig. 3a, Table S10). These genes include *TaWOX5* and *TaLBD13* involved in root development. Interestingly, some auxin transport and signaling pathway genes, such as *TaPIN1* and *TaARF10* are regulated by both B-ARR and ARF (Fig. 3e), suggesting a crosstalk between auxin and cytokinin signaling.

Thus, auxin signaling is important for initiation and driving the cell fate transition during callus formation and paves the way for later organ differentiation by activating and crosstalk with cytokinin signaling.

### Sequential transcriptional regulatory network in driving wheat shoot regeneration

The high coordination between transcription and chromatin accessibility suggests that cis-motif and trans-factor mediated transcriptional regulatory network would govern wheat shoot regeneration. To bridge TFs and target genes, we identified footprints from ATAC-seq data (Extended Data Fig. 4a). Most footprints are distributed in the middle of ATAC-seq peaks, about 10-50bp in width, and nearly half of them could match the TF binding motifs of *Arabidopsis* (Extended Data Fig. 4b, c, d). As expected, the sequence of footprints is more conserved than that in the intergenic regions (Extended Data Fig. 4e). We analyzed the differential footprints (Extended Data Fig. 4f, Table S11) and found that TF activity of known regulators involved in plant regeneration changed significantly, such as *LEC2, DOF5*.*6, WOX11* and *BBM* etc (Fig. 4a), suggesting potential functional importance. In addition, co-expression of TFs and target genes is also required (Extended Data Fig. 4a). Genes with similar expression patterns may be regulated by common upstream TFs. Therefore, we performed the enrichment analysis of upstream motifs of TFs for each RNA cluster (Fig. 4b, Table S12). Among these enriched motifs, there are some of known factors that can promote plant regeneration, such as *WOX11* and *WUS* in WOX family, *BBM* in AP2 family and *WIND3* in ERF family (Fig. 4b). Then the network was filtered according to the enrichment results, and a network containing 31,272 nodes and 8,856,326 edges mediated by 1,766 TFs was obtained (Extended Data Fig. 4g). Among them, 1,253 regulatory relationships between orthologous TFs and target genes in *Arabidopsis* have been verified in the yeast monohybrid system (Ikeuchi et al., 2019).

**Fig. 4.**
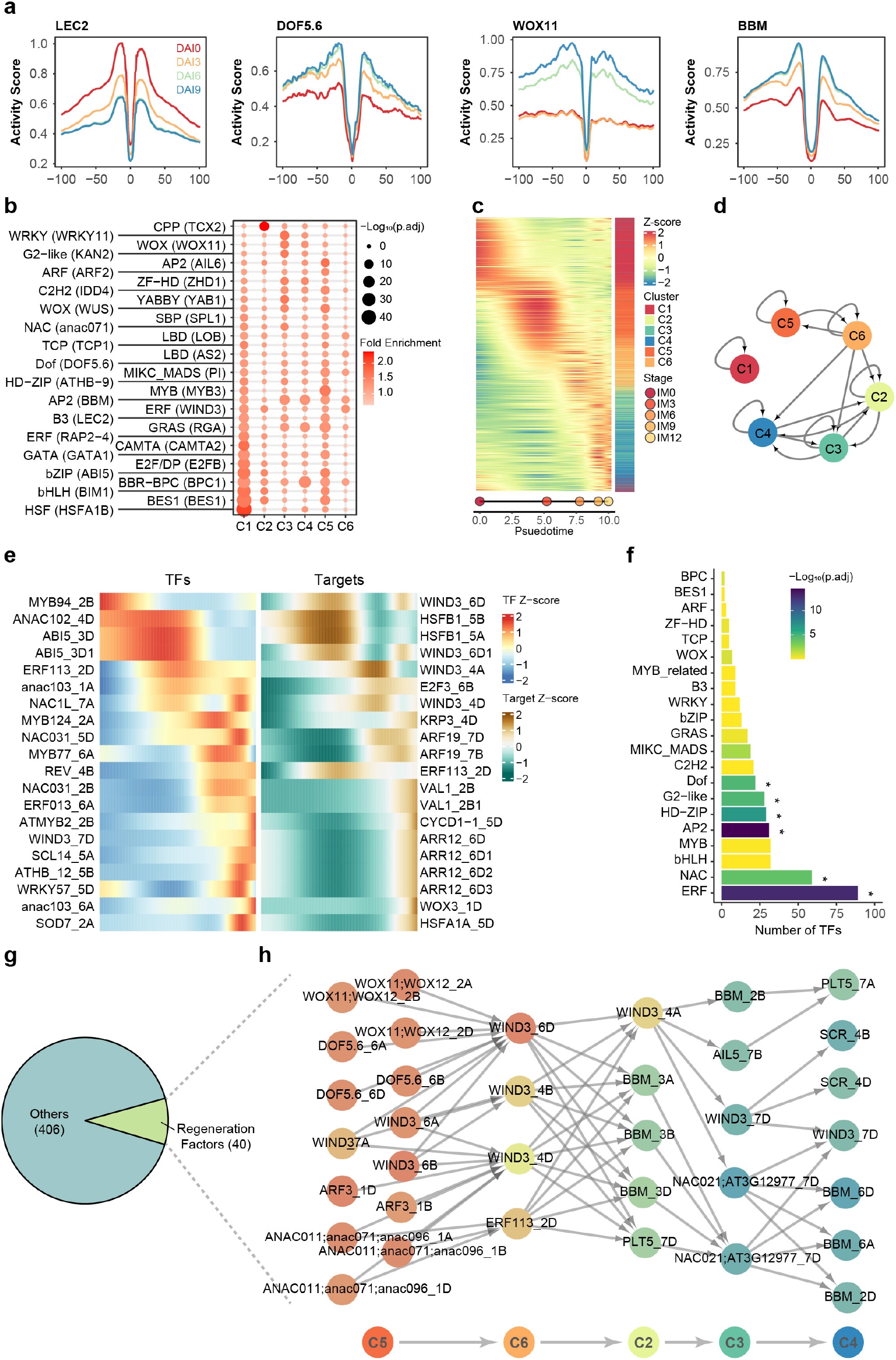
Transcriptional regulatory network governing wheat shoot regeneration. **a**, ATAC-seq footprint for LEC2, DOF5.6, WOX11 and BBM at different induction stages; Activity score reflects the chromatin accessibility. **b**, The enriched TF binding motifs for each RNA cluster of Fig. 1f. **c**, Gene sorted by time of peak expression during wheat shoot regeneration. **d**, Regulatory relationships between different RNA clusters (Fisher’s exact test, p.adj < 1e-6). **e**, Expression pattern of TFs and targets in the TRN. The regulatory relationship between TFs and targets has been demonstrated in *Arabidopsis*. **f**, The TF family enrichment analysis of core 446 TFs (Fisher’s exact test, ***: p.adj<0.001). **g**, The 446 core TFs contain 40 orthologous genes of regeneration factors in *Arabidopsis*. **h**, The regulatory network of orthologous genes of regeneration factors in *Arabidopsis*.

Pseudotime analysis was performed to determine the temporal order of gene expression. A clear sequential time-serial expression pattern of different clusters is observed, following the order of C1-C5-C6-C2-C3-C4 (Fig. 4c, Table S14). We further generated an intra-cluster regulatory network to explore the regulatory relationship between different clusters (Fig. 4d, Table S15). As expected, TFs in each cluster can regulate target genes within their own cluster. No regulatory relationship between C1, which specifically highly expressed at DAI0, and other clusters was observed, further emphasizing the effects of auxin treatment. Interestingly, we found a sequential regulatory relation among clusters C5-C6-C2-C3-C4, which in general follows the time line of gene expression (Fig. 4d). Therefore, we speculated that two different regulatory modes co-existing in driving wheat shoot regeneration (Extended Data Fig. 4h). Genes in each cluster are regulated by TFs in the same cluster (Type I) (Table S16), as well as genes in later expressed cluster are regulated by TFs in the previous expressed cluster (Type II) (Table S17). Indeed, we identified functionally related TFs in both regulatory modes, such *TaWIND3, TaDOF5*.*6, TaWOX5, TaBBM, TaAIL5* and *TaSCR* (Extended Data Fig. 4i, j). Generally, TFs and target genes show similar dynamic transcriptional profile during the regeneration process (Fig. 4e). We further extracted a total of 446 core TFs by combing the two types of regulation modes (Table S18). Among them, AP2, ERF, HD-ZIP, DOF, G2-like and NAC family are highly enriched (Fig. 4f) and 40 TFs are the orthologous genes of regeneration factors functionally tested in *Arabidopsis* (Fig. 4g). Interestingly, those genes are sequentially regulated during wheat shoot regeneration (Fig. 4h, Table S19). For instance, *TaWOX11, TaDOF5*.*6* and *TaWIND3* are first activated, and then they activate *TaBBM, TaAIL5* and *TaSCR* to promote the callus formation (Fig. 4h).

In summary, we used time course transcriptome and chromatin accessibility data to construct a temporal TRN, which reflects the continuity of the wheat shoot regeneration process. The TRN contains hundreds of core TFs and captures the regulatory relationships among known regeneration factors in model plant species.

### Early callus induction factors are distinct between wheat and *Arabidopsis thaliana*

Since the core TRN driving regeneration contains conserved TFs in both *Arabidopsis* and wheat, we therefore further compared the process in these two species (Raw data for *Arabidopsis* regeneration is from Wu et al., 2022). We performed differential expression analysis between stages within each species and identified orthologous genes from DEGs (Table S20, Table S21). We noticed that most DEGs have orthologous genes, while only 35% are species-specific (Fig. 5a). For comparison, we focused on the orthologous genes of DEGs (Group III). Hierarchical clustering and principal component analysis were performed using z-score normalized gene expression and chromatin accessibility profiles within species (Fig. 5b, Extended Data Fig. 5a). The expression peaks of *WOX12* in *Arabidopsis* and wheat were observed at CIM3d and DAI3, respectively, indicating the root founder cell stage (Fig. 3a, Extended Data Fig. 5b). From *WOX5* expression pattern, CIM5d-7d and DAI6-9-12 were presumed to be the root meristem stage (Fig.3a, Extended Data Fig. 5b). Overall, the callus induction process of wheat and *Arabidopsis* is conserved, reflected by the fact that samples at the same induction stage are more similar between the two species than different induction stages within the same species (Fig. 5b). However, some regeneration factors have different expression pattern in wheat and *Arabidopsis* (Fig. 5c). In addition, the motif activities of certain TFs are also different between wheat and *Arabidopsis* (Extended Data Fig. 5c, Table S22). Thus, though orthologous genes are involved in the regeneration process of both wheat and *Arabidopsis*, species-specific expression pattern and transcriptional regulatory route may exist.

**Fig. 5.**
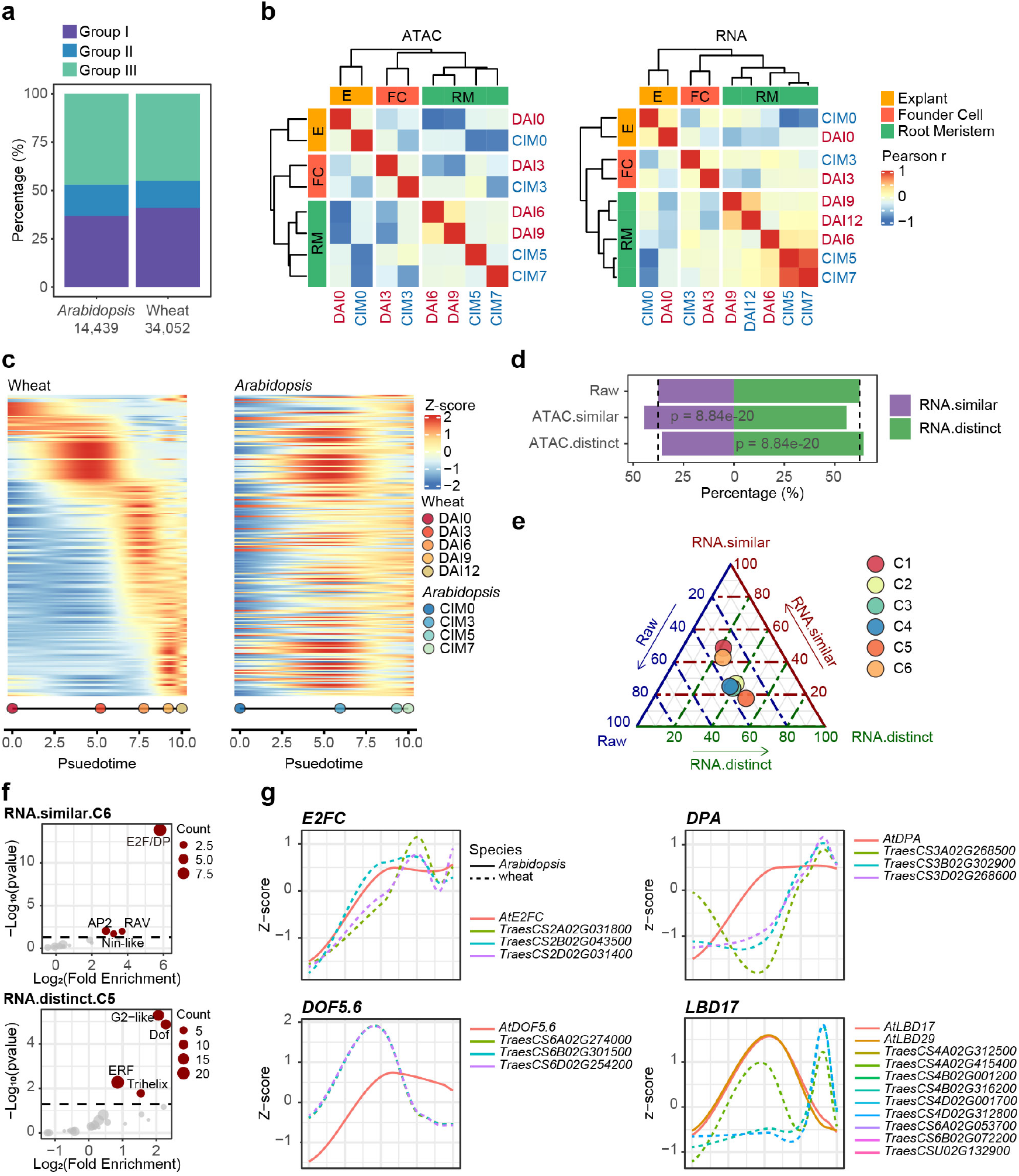
Comparative analysis of regeneration process between wheat and Arabidopsis. **a**, Classification of DEGs in wheat and *Arabidopsis*; Group I represents species specific DEGs; Group II represents DEGs whose orthologous genes in another species are not expressed or not differentially expressed; Group III represents DEGs whose orthologous genes in another species are also differentially expressed. **b**, Hierarchical clustering analysis of RNA-seq and ATAC-seq during regeneration in wheat and *Arabidopsis*; E: Explant; FC: Founder cell; RM: Root meristem; The red font represents samples from wheat and the blue font represents samples from *Arabidopsis*. **c**, The sorted expression pattern of regeneration factor in wheat and *Arabidopsis*. **d**, The proportion of genes with similar and distinct expression pattern during regeneration in wheat and *Arabidopsis*. **e**, Over representation analysis of genes with similar and distinct expression pattern in wheat RNA clusters. **f**, TF family enrichment analysis of genes with similar expression pattern in C6 and genes with distinct expression pattern in C5. **g**, The expression pattern of *E2FC, DPA, DOF5*.*6* and *LBD17* in wheat and *Arabidopsis*.

Based on the correlation of expression patterns, orthologous genes were divided into two categories: genes with similar expression pattern or distinct expression pattern (Table S20). The similarity or difference of gene expression pattern is highly related to the chromatin accessibility in the proximal of orthologous genes (Fig. 5d). To explore the characteristics of these two types of genes, we performed an over representation analysis in the RNA cluster (Fig. 5e, Table S23). We found that orthologous genes with similar expression pattern are mostly enriched in C1 and C6, which cover genes specifically highly expressed in the explant before auxin addition (C1) and genes generally induced by auxin treatment (C6) (Fig. 5e). This again speaks of the similarity of callus induction process between wheat and *Arabidopsis*. While orthologous genes with distinct expression pattern are enriched in C5, which is transiently induced by auxin at early callus induction stage (Fig. 5e). Genes in C1 and C6 with similar expression pattern are enriched in ERF and E2F family (Extended Data Fig. 5e, Fig. 5f, Table S24), including *TaE2FC* and *DP-LIKE PROTEIN A* (*TaDPA)* (Fig. 5g), while genes in C5 with distinct expression pattern are enriched in DOF and G2-like family (Fig. 5f, Table S24), such as *TaDOF5*.*6* (Fig. 5g). In addition, the activity score of *DOF5*.*6* footprint is consistently up-regulated during callus induction in wheat (Fig. 4a), but shows a different pattern in *Arabidopsis* (Extended Data Fig. 5f). For example, the expression level of *DOF5*.*6* is up regulated at the early stage of callus induction in wheat but at the later stage in *Arabidopsis* (Fig. 5g).

As the orthologue genes with distinct expression pattern are enriched in C5, we wonder whether early stage of callus induction is species-specific. Therefore, we performed k-means cluster analysis of DEGs in *Arabidopsis* and identified cluster A3 with similar pattern to wheat cluster C5 (Extended Data Fig. 5d, Table S21). Different from DOF family enriched in wheat cluster C5, LBD and NAC families are enriched in *Arabidopsis* cluster A3 (Extended Data Fig. 5g, Table S24). In addition, the motif activity of LBD was significantly increase in CIM3d of *Arabidopsis*, but not in wheat until DAI6 (Extended Data Fig. 5h), consistent with its expression pattern in both species (Fig. 5g). Interestingly, overexpression of LBD was reported to promote the regeneration process in *Arabidopsis* (Fan et al., 2012).

In short, we found that the callus induction process is generally conserved between wheat and *Arabidopsis*. However, there is species-specific genes expression and transcriptional regulation modules at early callus induction stage.

### DOF TFs boost regeneration efficiency of wheat varieties

As species-specific enriched TFs at early callus induction in wheat, we wonder whether manipulation of DOF family TFs expression could improve the regeneration efficiency in wheat. Among the DOF TFs in C5, the expression level of *TaDOF5*.*6* and *TaDOF3*.*4* are most sharply up-regulated upon auxin treatment (DAI3) (Extended Data Fig. 6a), along with the removal of H3K27me3 (Fig. 6a). Meanwhile, chromatin accessibility of *TaDOF5*.*6* and *TaDOF3*.*4* binding sites is significantly increased at DAI3 and remains high at DAI6 and DAI9 (Fig. 4a, Extended Data Fig. 6b). In addition, *TaDOF5*.*6* and *TaDOF3*.*4* occupy the early transcriptional hierarchy within the core TRN, which can regulate genes related to cell cycle, cell proliferation and lateral root development, such as *TaBBM, TaE2F3, TaARR10* and *TaLBD17* (Extended Data Fig. 6c, d, Table S25). Thus, we choose *TaDOF5*.*6* and *TaDOF3*.*4* as potential ‘boosters’ for testing effects on regeneration efficiency.

**Fig. 6.**
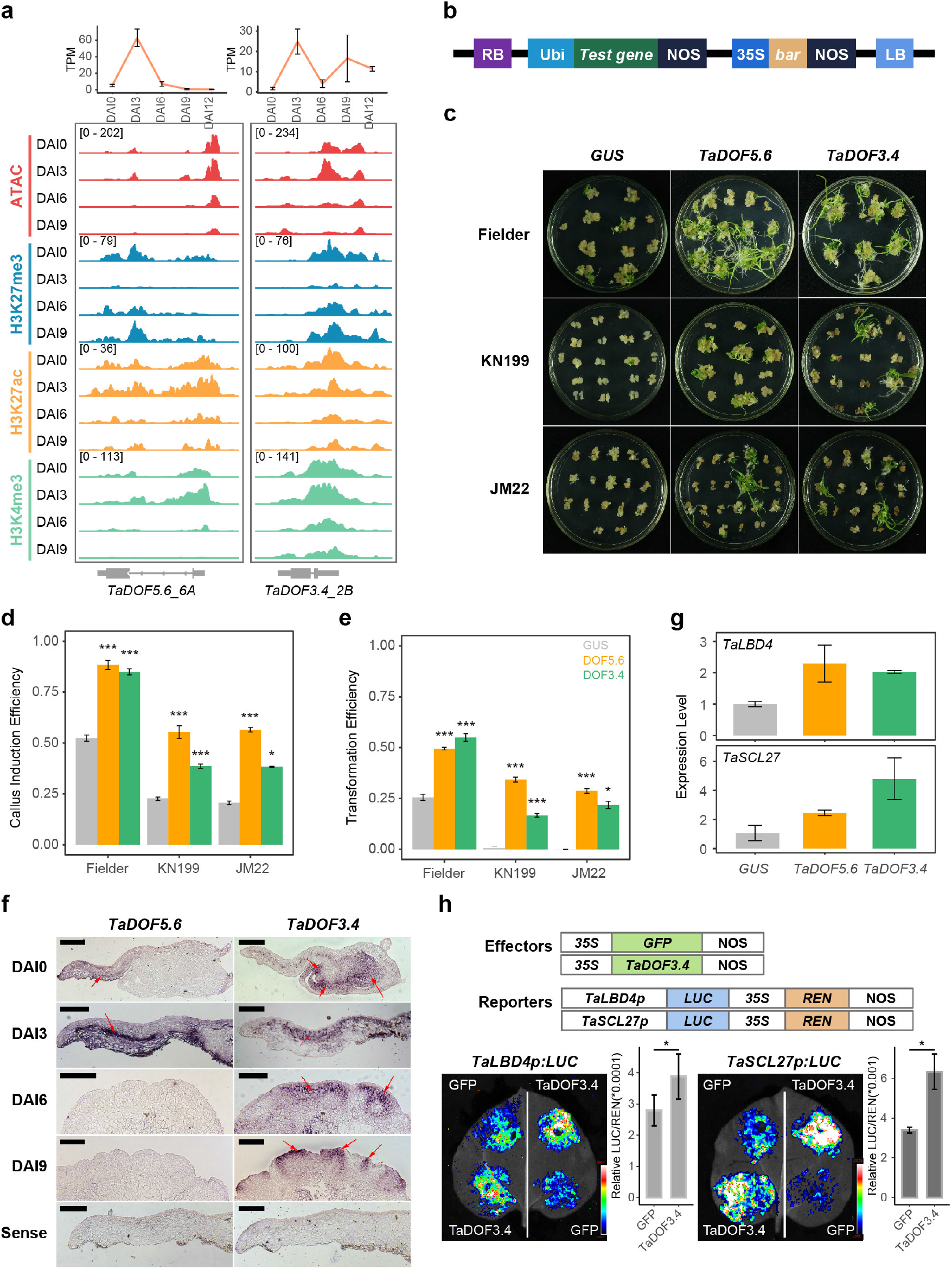
TaDOF5.6 and TaDOF3.4 boost wheat genetic transformation efficiency. **a**, The dynamic transcription and epigenetic modification track for *TaDOF5*.*6_6A* and *TaDOF3*.*4_2B* during regeneration. **b**, Structure of the over-expression vector pc186; RB: Right boundary; Ubi: Maize ubiquitin gene promoter; NOS: Carmine synthase gene terminator; 35S: Cauliflower mosaic virus 35S promoter; *bar*: Phosphinothricm acetyltransferase gene; LB: left boundary. **c**, Shoot regeneration of Fielder, KN199 and JM22 transformed with pc186 vectors containing either *GUS, TaDOF5*.*6* or *TaDOF3*.*4*. **d**,**e**, Callus induction efficiency (**d**) and transformation efficiency (**e**) of Fielder, KN199 and JM22 transformed with vectors containing either *GUS, TaDOF5*.*6* or *TaDOF3*.*4* (Student’s t-test; *: p<0.05; ** p<0.01; ***: p<0.001). **f**, *In situ* hybridization of *TaDOF5*.*6* and *TaDOF3*.*4* during wheat shoot regeneration; The arrows indicate where the gene is highly expressed. **g**, The expression pattern of *TaLBD4* and *TaSCL27* in DAI9 of transformed with *GUS, TaDOF5*.*6* or *TaDOF3*.*4*. **h**, Luciferase reporter assays validation of transcriptional activation capability of *TaDOF3*.*4* on targets *TaLBD4* and *TaSCL27* (Student’s t-test; *: p<0.05).

First, we co-transformed either *TaDOF5*.*6* or *TaDOF3*.*4* for regeneration from immature embryo of wheat variety Fielder (Fig. 6b). As compared to *GUS* negative control, both *TaDOF5*.*6* and *TaDOF3*.*4* co-transformation could significantly increase the callus induction efficiency and transformation efficiency (Fig. 6c, Table S26), from 52% to 88% and 85% for callus induction efficiency, and 26% to 50% and 55% for transformation efficiency, respectively (Fig. 6d, e, Table S45). Second, since Fielder is easy for regeneration, we expanded our test to some wheat varieties that is hard for regeneration, such as Jimai 22 (JM22) and KN199 (Wang et al., 2017; Zhang et al., 2018). Of note, compared with negative control, which shows low or even no regeneration in JM22 and KN199 variety, co-transform of *TaDOF5*.*6* and *TaDOF3*.*4* could increase the transformation efficiency of JM22 to 29% and 22%, and the transformation efficiency of KN199 to 34% and 17%, respectively (Fig. 6d, e, Table S26). Thus, *TaDOF5*.*6* and *TaDOF3*.*4* could elevate wheat shoot regeneration efficiency from a broad spectrum of wheat varieties.

Based on the TRN, we speculated that *TaDOF5*.*6* and *TaDOF3*.*4* may directly regulate the expression of genes related to root and shoot meristem formation, such as *LBDs* (Fan et al., 2012), *YABs* (Gordon et al., 2007) and *SCARECROW-LIKE 27* (*SCL27)* (Schulze et al., 2010) (Extended Data Fig. 6c, d) to boost the regeneration efficiency of wheat. *In situ* hybridization confirmed that *TaDOF5*.*6* is expressed only in the procambial area at DAI3, whereas *TaDOF3*.*4* id expressed not only in the three to four layers below the epidermis at DAI3, but also in embryogenic cells at DAI6 and promeristem of shoot primordia at DAI9 (Fig. 6f). We next sampled tissues from DAI9 of co-transformed *TaDOF5*.*6* and *TaDOF3*.*4* and *GUS* control. Indeed, *TaLBD4* and *TaSCL27* are higher expression in *TaDOF3*.*4* and *TaDOF5*.*6* co-transformed tissue in compared with *GUS* (Fig. 6g). Transcriptional reporter assay further validated the regulation of *TaDOF3*.*4* to *TaLBD4* and *TaSCL27* (Fig. 6h).

Thus, *TaDOF5*.*6* and *TaDOF3*.*4* serve as novel boosters for increasing wheat shoot regeneration efficiency via promoting root and shoot meristem development.

## Discussion

The success of genetic transformation is largely dependent on the regeneration capacity of explant (Hu et al., 2017). Hierarchy transcriptional regulation driven by hormones signaling in the context of chromatin dynamics for regeneration have been well studied in *Arabidopsis* and numerous genes that regulate regeneration have been identified and applied in crops for transformation efficiency improvement (Ikeuchi et al., 2019; Wang et al., 2020a; Wu et al., 2022). However, we know very little about the regeneration process of monocot crops, such as wheat. Therefore, we generated the first set of transcriptome and epigenome data during wheat shoot regeneration, which provides valuable data resources for systematic study of molecular insights for regeneration in monocot crops and mining of regeneration efficiency accelerators (Fig.1).

### Epigenetic layer regulation affects wheat shoot regeneration efficiency

Gene expression during wheat shoot regeneration is sequential and co-regulated by chromatin accessibility, H3K27me3 and H3K4me3 (Fig. 2, Fig. 7). Gain of chromatin accessibility is along with activation of key genes that mediating cell fate transition during the process of callus induction (Fig. 2, Fig. 7). One would speculate that accessible chromatin environments may increase the chance for regeneration. This is evidenced in *Arabidopsis* for regeneration from protoplast, which introducing randomly opened chromatin regions during the protoplast generating process (Xu et al., 2021). As well, increase the histone acetylation, which could increase chromatin accessibility, by chemical treatment such as Trichostatin A or sodium butyrate could promote the regeneration process in wheat (Bie et al., 2020; Jiang et al., 2017).

**Fig. 7.**
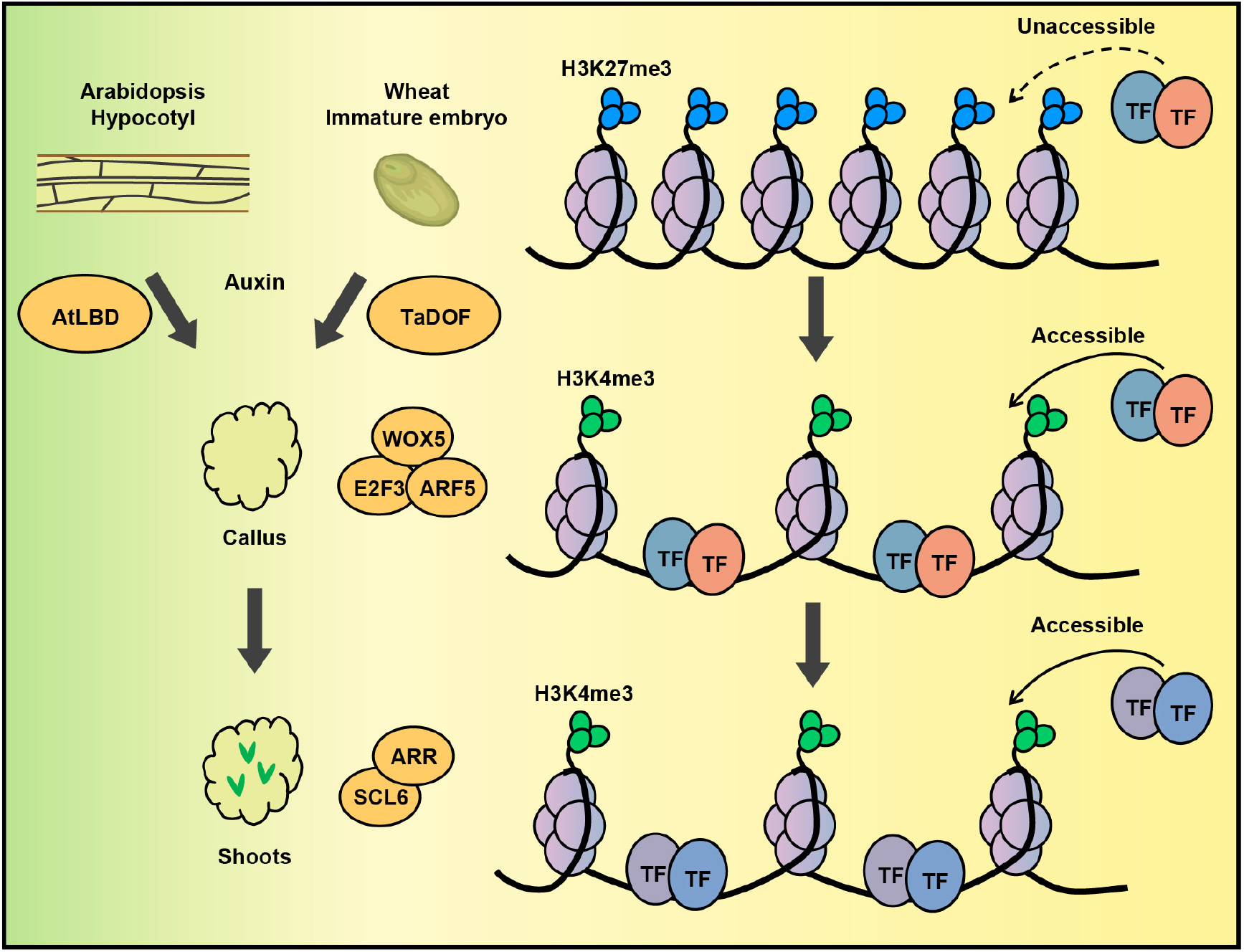
The working model for epigenetic associated transcriptional regulation of wheat shoot regeneration process.

In addition, we also found that H3K27me3 decrease and H3K4me3 increase at genes activated in the late stage of callus induction in wheat (Fig. 2, Fig. 7). Most of these genes are associated with auxin signaling and root development, such as *TaPIN1* and *TaLBD17*, suggesting that the removal of H3K27me3 and the increase of H3K4me3 are important for wheat shoot regeneration. Similarly, H3K27me3 and H3K4me3 are involved in callus formation in rice (Zhao et al., 2020). In *Arabidopsis*, the reprogramming of H3K27me3 is critical for the acquisition of totipotency, as callus formation is impaired in H3K27me3 methyltransferase *curly leaf swinger* (*clf swn*) mutant (He et al., 2012). In addition, the H3K4me3 methyltransferase *ATX4* could promote shoot regeneration (Lee et al., 2019). Therefore, it will be worth to test the effect of H3K27me3 demethylase and H3K4me3 methyltransferase on wheat shoot regeneration.

### Identification of boosters for wheat shoot regeneration by comparison study

Callus formation occurs regardless of species and explants induced on CIM, but the detailed process of monocots and dicots may different. By comparing the regeneration process of wheat and *Arabidopsis*, we found that the callus induction process is generally conserved, especially the genes related to cell proliferation. However, in the early stage of callus formation, the earliest up-regulated TFs were different between the two species, namely, the DOF and G2-like families in wheat, while LBD and NAC families in *Arabidopsis* (Fig. 5, Fig. 7). However, we cannot tell whether this difference is due to explants diversity or species related.

Interestingly, the LBD and NAC TFs were proved to accelerate regeneration in *Arabidopsis* (Daimon et al., 2003; Fan et al., 2012), suggesting the early responsive factors for auxin may play important role in regeneration. Following this clue, we tested DOF family members for the effects on wheat shoot regeneration. Indeed, both *TaDOF5*.*6* and *TaDOF3*.*4* could boost the regeneration efficiency of Fielder and other varieties that hard to regenerate, such as JM22 and KN199 (Fig. 6). Thus, such comparison study could provide species-specific candidates for boosting the regeneration efficiency. Future work on that wheat specific TFs may give more options for manipulation of wheat transformation efficiency.

### TRN facilitates functional study and may mediate varieties difference

Based on the high positive correlation between chromatin accessibility and transcription, we generated a time-course hierarchy transcriptional regulatory network that drives wheat shoot regeneration via integrating of RNA-seq and ATAC-seq data (Fig. 4). A total of 446 core TFs were identified including 40 orthologous genes of known regeneration factors in *Arabidopsis* (Fig. 4). Certain factors of the 446 TFs are expressed at different stages, indicating a potential stage-specific role. Thus, integrating multiple stage-specific factors may further improve the transformation efficiency as compared to individual booster. The hierarchy TRN also extends the understanding of molecular regulation mechanism of wheat shoot regeneration process, and facilitate gene functional study for individual factors. For instance, the TRN provides the downstream targets that regulated by DOF TFs for accelerating the regeneration efficiency, such as root and shoot meristem related genes *TaLBD4* and *TaSCL27* (Fig. 6).

Of note, the core TRN was constructed in the easiest transformed wheat variety Fielder background, which may provide a candidate list for mediating the varied regeneration efficiency among different wheat varieties. To support this view, we further investigated the genetic variations (Hao et al., 2020; Wang et al., 2020b) in the coding region and open chromatin region of the 446 core TFs in 12 wheat varieties with different regeneration efficiency range (Zhang et al., 2018). Among them, 247 genes have coding region variations and 188 genes have regulatory region variations. In particular, genetic variations of three TFs in WOX, C2H2 and NAC family are correlated with differentiated callus rate (Table S27). Further analysis would focus on those factors for understanding the genotype-dependent regeneration difference in wheat varieties.

In summary, we integrated multi-omics data to reveal the transcriptional regulatory network and epigenetic dynamic during wheat shoot regeneration, and identified two novel genes that can improve wheat transformation efficiency. Furthermore, since callus is not a homogeneous mass of cells, but rather resembles a root primordium in which the middle layer of cells is necessary for regeneration (Atta et al., 2009), omics data at single cell level may provide finer clues for studies of crop regeneration (Liu et al., 2022).

## Methods

### Plant materials and growth conditions

Hexaploid wheat varieties Fielder, KN199 and JM22 were used in the genetic transformation. Fielder was kindly provided by Japan Tobacco Inc. Fielder seeds (spring wheat) were directly sown in pots (21 cm height × 23 cm diameter), while KN199 and JM22 were grown in 6° C incubator for 35 days after germinated 2 days. All wheat genotypes were maintained with a growth condition regime of 24°C, 16 h light/18°C, 8 h dark with 300 μmol/m^2^/s light intensity at 40% humidity in climate chamber.

### Plant materials used for RNA-seq, ATAC-seq and CUT&Tag

The immature embryos of Fielder were cultured generally following the described protocol (Ishida et al., 2015) with slight modifications. In short, immature embryos were isolated from the immature seeds into WLS liquid medium (1/10 Murashige and Skoog (MS) salts, 1/10 Murashige and Skoog (MS) vitamins, glucose 10 g/L, 2-(N-morpholino) ethanesulfonic acid (MES) 0.5 g/L, pH 5.8) at room temperature, and incubated on WLS-As medium (WLS liquid medium plus AgNO_3_ 0.85 mg/L, CuSO_4_ 5H_2_O 1.25 mg/L, acetosyringone 100 μM and agarose 8 g/L) at 23°C in the dark for 2 days. After cocultivation, embryonic axes were excised with scalpel and forceps, and remaining scutella were transferred to WLS-Res medium (MS salts, MS vitamins, glutamine 0.5 g/L, casein hydrolysate 0.1 g/L, MgCl_2_ 6H_2_O 0.75 g/L, maltose 40 g/L, 2,4-D 0.5 mg/L, picloram 2.2 mg/L, AgNO_3_ 0.85 mg/L, ascorbic acid 100 mg/L, MES 1.95 g/L and agarose 5 g/L) at 25°C in the dark for 19 days. Each callus was cut into two pieces with scalpel and transferred to WLS-Res medium for 3 weeks. Proliferated calli were differentiated on LSZ medium (MS salts, modified LS vitamins, zeatin 5 mg/L, sucrose 20 g/L, CuSO_4_ 5H_2_O 2.5 mg/L, MES 0.5 g/L and phytagel 3 g/L) for regeneration culture at 25°C with 100 μmol/m2/s light for 10 days. Regenerated plants were transferred on the LSF medium (MS salts, modified LS vitamins, IBA 0.2mg/L, sucrose 15 g/L, MES 0.5 g/L and phytagel 3 g/L) for elongation and root formation.

Materials at 0 3, 6, 9 and 12 DAI were photographed with JSM-6610LV scanning electron microscope (JEOL, Japan) for morphological observation. According to the above observation, wheat materials for RNA-seq were collected on the surface of callus at DAI3, DAI6, DAI9 and DAI12, while ATAC-seq and CUT&Tag at DAI3, DAI6 and DAI9, respectively. It is important to note that material at DAI0 was scutellum of immature embryo after excision of embryo axis without tissue culture. Two or three biological replicates were performed. For each biological replicate, approximately 30-60 immature embryo explants cultured were collected at different tissue culture stages.

### RNA-seq, ATAC-seq and CUT&Tag experiment

Total RNAs were extracted using Ultrapure RNA Kit (CWBIO, Cat No. CW0581M, China) according to the manufacturer’s instruction. The products were sequenced using an Illumina NovaSeq 6000 by Gene Denovo Biotechnology Co. (Guangzhou, China) and RNA-seq data were obtained from three biological replicates.

ATAC-seq and CUT&Tag experiment were done follow the previous described method (Zhao et al., 2022). Tn5 transposase used and tagmentation assay is done following the manual (Vazyme Biotech, TD501-01). Libraries were purified with AMPure beads (Beckman, A63881) and sequenced using the Illumina Novaseq platform at Annoroad Gene Technology. Antibodies used for histone modifications are the same as previous reported (Zhao et al., 2022)

### Plasmid construction and wheat transformation

Full length cDNA of *TaDOF3*.*4* or *TaDOF5*.*6* was cloned into pENTR™/D-TOPO® (ThermoFisher Scientific, Cat No. K2400-20SP) and inserted into the Gateway-compatible over-expression vector pc186 after sequenced. The control vector of GUS was constructed using the above method. All constructed vectors were transformed into callus induced from immature embryo via Agrobacterium-mediated transformation (strain EHA105) following the described protocol (Ishida et al., 2015) except for MS salts instead of LS salts in all media. The pc186 vector was kindly provided by Dr. Daolin Fu at College of Agronomy, Shandong Agricultural University, Tai’an, Shandong, China. Strain EHA105 was kindly provided by Japan Tobacco Inc.

### Transformation efficiency and statistical analysis

Explants, callus, and callus with at least one positive plantlet were counted after cultured on LSF medium for 10 days. Experimental data were analyzed with Student’s t-test t in R. Calculation methods as follows:

Callus induction efficiency (%): the number of callus / the number of explants incubated × 100%.

Transformation efficiency (%): the number of callus with at least one shoot / the number of explants incubated × 100%.

### *In situ* hybridization

*In situ* hybridization and detection of hybridized signals were carried out as described by previous study (Meng et al., 2017). The immature embryo of 14 days after anthesis and callus with tissue cultured at DAI3, DAI6 and DAI9 was fixed in 0.1 M phosphate buffer (pH 7.0) and 4% v/v paraformaldehyde (Sigma, USA) overnight at 4°C. After dehydration with graded ethanol and vitrification by dimethylbenzene, the specimens were embedded in paraplast (Sigma, USA) and sectioned at 8.0 μm. Antisense and sense RNA probes of *TaDOF5*.*6* and *TaDOF3*.*4* genes were synthesized using a DIG RNA Labeling Kit (SP6/T7) (Roche, Cat No. 11175025910, USA). The oligonucleotide primers for *in situ* hybridization are listed in Table S28.

### Quantitively RT-PCR analysis

Total RNA was extracted using Ultrapure RNA Kit (CWBIO, Cat No. CW0581M, China). First-strand cDNA was synthesized from 2 mg of DNase I-treated total RNA using the TransScript First Strand cDNA Synthesis SuperMix Kit (TransGen) as recommended by the manufacturer and was stored at -20°C. Quantitative PCR was performed using the ChamQ Universal SYBR qPCR Master Mix (Q711-02, Vazyme Co., Nanjing, China) by QuantStudio5 appliedbiosystems. Samples were normalized with *TaTublin* expression; relative expression levels were measured using the 2ΔCtanalysis method. Primer sequences were listed in Table S28.

### Luciferase reporter assay

The luciferase reporter assay was done follow the described method (Zhao et al., 2022). The genomic sequence of down-stream targets *TaLBD4* and *TaSCL27* promoter containing the DOF binding site were amplified and fused in-frame with the CP461-LUC vector digested with EcoRI to generate the reporter construct target-pro: LUC. The CDS of up-stream TF *TaDOF3*.*4* was cloned into the SpeI digested PTF101 vector to generate the effector construct 35Spro: TF. Primer sequences were listed in Table S28.

### RNA-seq data analysis

Raw reads were processed by fastp (v0.23.1) (Chen et al., 2018) with “--detect_adapter_for_pe” parameter to remove adapters, trim low quality bases and filter bad reads. Clean reads were aligned to wheat reference genome (IWGSC RefSeq v1.0(International Wheat Genome Sequencing Consortium (IWGSC), 2018), downloaded from https://urgi.versailles.inra.fr/download/iwgsc/IWGSC_RefSeq_Assemblies/v1.0/) using hisat2 (v2.0.5) (Kim et al., 2019)and the aligned reads were sorted and converted into bam files using samtools (v1.5) (Danecek et al., 2021). featureCounts (v2.0.1) (Liao et al., 2014) was used to count the number of paired mapping reads that overlap each annotated gene (IWGSC Annotation v1.1) with the parameter “-p -P -B -C”. The counts matrixs were used as inputs to identify differentially expressed genes using R package DESeq2 (v1.28.1) (Love et al., 2014), with a threshold of p.adj<0.05 and abs(log2FoldChange)>1. TPM (Transcripts Per Kilobase Million) values generated from the counts matrix were used to characterize gene expression and used for PCA analysis and hierarchical clustering analysis. K-means cluster analysis of differentially expressed genes were performed in R (v4.0.2) using Z-scaled TPM and the gene expression heatmap was displayed using R package ComplexHeatmap (v2.4.3) (Gu et al., 2016). The gene ontology enrichment analysis was done on the GOC website (http://geneontology.org/) which is supported by PANTHER (Mi et al., 2019). The identification and family classification of wheat TFs were predicted on the PlantTFdb website (http://planttfdb.gao-lab.org/) (Tian et al., 2020). Enrichment analysis of TF families for DEGs was achieved using the enricher function in R package clusterProlifer (v3.16.1) (Wu et al., 2021).

### CUT&Tag and ATAC-seq data analysis

Raw reads were processed by fastq (v0.23.1) with “--detect_adapter_for_pe” parameter to remove adapters, trim low quality bases and filter bad reads. Clean reads were aligned to wheat reference genome (IWGSC RefSeq v1.0) using bwa mem algorithm (v 0.7.17) (H and R, 2009) with “-M” parameter. The aligned reads were sorted and filtered by samtools (v1.5) with “-F 1804 -f 2 -q 30” parameter. Picard (v 2.23.3) was used to removed the duplicated reads. The de-duplicated bam files from two biological replicates were merged by samtools (v1.5) and converted into bigwig files using bamCoverage provided by deeptools (v3.5.0) (Ramírez et al., 2016) with parameters “-bs 10 --effectiveGenomeSize 14600000000 --normalizeUsing RPKM --smoothLength 50”. The bigwig files were visualized using deeptools (v3.5.0) and IGV (v2.8.0.01) (Thorvaldsdóttir et al., 2013).

The peak calling was done using merged bam files by macs2 (v 2.2.7.1) (Zhang et al., 2008). For ATAC-seq data, the parameter of peak calling using macs2 is “-q 0.05 -f BAMPE --nomodel --extsize 200 --shift -100 -g 14600000000”. For narrow peak of H3K27ac and H3K4me3, the peak calling parameter is “-p 1e-3 -f BAMPE -g 14600000000 --keep-dup all”. For broad peak of H3K27me3, the peak calling parameter is “-f BAMPE -g 14600000000 --keep-dup all --broad --broad-cutoff 0.05”. Peaks from all samples of the same data type were merged using bedtools (v2.29.2) (Quinlan and Hall, 2010) to generate the reference peaks. Peak was annotated to the wheat genome using the R package ChIPseeker (v1.24.0) (Yu et al., 2015). The whole genome was divided into three regions: promoter (from -3000bp to +500bp of TSS), genic (from +500bp of TSS to TES) and distal (other). Peaks located in promoter and genic regions were annotated to genes that overlap with them, while peaks in distal region were annotated to nearest gene/TSS.

For quantification of ATAC-seq and CUT&Tag data, reads count under reference peak and the normalized CPM (DBA_SCORE_TMM_READS_EFFECTIVE_CPM) values were generated using R package DiffBind (v2.16.2). The CPM values of reference peak were used to do PCA analysis and hierarchical clustering analysis. The raw peak counts were used as input to identify differentially accessible peaks and differentially marked peaks by histone modification using R package DESeq2 (v1.28.1). For differentially peaks located in promoter and genic region, the CPM values was z-scaled to perform k-means clustering analysis in R. The heatmaps centered on peak were created using computeMatrix and plotHeatmap from deeptools (v3.5.0).

To calculate correlations between transcriptome and epigenome data, z-scaled TPM values averaged over all DEGs within each RNA cluster and z-scaled CPM values averaged over peaks annotated to promoter and genic regions of these DEGs were calculated. Then pearson correlation coefficients were calculated in R for each RNA cluster.

For each RNA cluster, the over representation analysis in open chromatin and histone modification cluster was performed using the enricher function in R package clusterProlifer (v3.16.1).

### Identification of ARF and type-B ARR target genes

The target genes of ARF should meet two criteria: Firstly, ATAC-seq peak in gene promoter or genic region contains ARF binding motif; Secondly, the genes belong to any cluster of C2, C3, C4, C5 and C6.

The target genes of type-B ARR should meet two criteria: Firstly, ATAC-seq peak in gene promoter or genic region contains type-B ARR binding motif; Secondly, the genes belong to any cluster of C2, C3 and C4. The binding motif of ARFs and type-B ARRs in *Arabidopsis* were downloaded from PlantTFdb. Fimo (v5.4.1) (Grant et al., 2011) was used to scan for motifs within peaks.

### ATAC-seq footprint analysis

HINT (v0.13.1) (Li et al., 2019) was used to identify footprints for ATAC-seq and the *Arabidopsis* motifs from PlantTFdb was used as motifs set. Based on the width of footprints, sequence conservation analysis was performed by randomly dissecting one million 25bp sequences in footprints, exons and intergenic regions. Sequence variation data of hexaploid wheat was obtained from public publications (Zhou et al., 2020).

### Pseudotime analysis

Pseudotime analysis was performed as previously described (Leiboff and Hake, 2019) with some modifications. All expressed genes (TPM > 1 in any sample) were used to separate samples by PCA. Then, the euclidean distance of adjacent samples along the development trajectory were calculated to produce developmental time units (DTU) values, scaled from 0.0 to 10.0. For each gens, the z-scaled expression values was interpolated into 500 points along its pseudotime metric using the loess function in R to allow for smooth, continuous comparisons. To generate phasigrams (Levin et al., 2016), PCA on z-scaled expression values was performed for each gene. Component 1 and component 2 formed a circle on the PCA plot. Genes were ordered based on their time of peak expression using Atan2 function in R, which was visualized in Fig. 4c by ComplexHeatmap (v2.4.3).

### TRNs construction

The network was set up in a similar way as previously reported (Hoang et al., 2020). To build the core regulatory networks, we only focus on DEGs with TPM values higher than 1 in any samples and ATAC-seq peaks located in the promoter or genic region of these genes. Firstly, HINT (v0.13.1) was used to identify the footprints within ATAC-seq peaks and the motifs within the footprints. The motif information was obtained from all *Arabidopsis* TFs in PlantTFdb. In this way, we can obtain the regulatory relationship between motifs and target genes. Secondly, to match the motifs of *Arabidopsis* TFs with those of wheat, we used PlantTFdb to predict the TFs and their families in wheat. TFs in wheat are mapped to *Arabidopsis* TFs only if they meet the following two criteria: 1) In blastp (v 2.10.1) (Camacho et al., 2009) results, evalue<1e-5 and identity > 30%; 2) TFs of two species belong to the same family. In this way, the regulatory relationships between wheat TFs and target genes were obtained through the TF binding motifs of *Arabidopsis*. Thirdly, considering that TFs and target genes need to be expressed simultaneously, we filtered the regulatory relationships according to the following principle: the TPM value of TFs must be higher than 1 when their target genes were highly expressed. For target genes of C1, C2, C3, C4, C5 and C6, the TPM value of their TFs must be higher than 1 in DAI0, DAI6, DAI9, DAI12, DAI3 and DAI3, respectively. Finally, we focus on those regulatory relationships linked by motifs enriched upstream of RNA clusters. Enrichment analysis was done using the R package clusterProlifer (v3.16.1) with a threshold p.adj < 1e-6. Therefore, we finally obtained a transcriptional regulatory network consisting of 1,766 TFs with 31,272 nodes and 8,856,326 edges.

Regulatory network between clusters were generated according to previously reported methods (Hoang et al., 2020). We used fisher test to calculate the significance of regulations from cluster C1 to cluster C2.

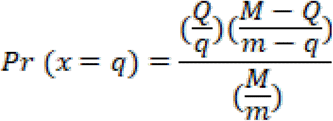

where *M* represents the total number of regulations, *Q* represents the total number of regulations from TFs in cluster C1, *m* represents the total number of regulations targeting cluster C2, *q* represents the number of regulations from cluster C1 to cluster C2, and the general form 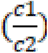 is a binomial coefficient. Pvalue < 1e-6 was set as the threshold to obtain regulatory network between clusters.

### RNA-seq and ATAC-seq analysis of *Arabidopsis*

The raw data of *Arabidopsis* RNA-seq and ATAC-seq were download from a previous publication (Wu et al., 2022). The RNA-seq and ATAC-seq data in *Arabidopsis* were analyzed in the same manner as in wheat, except that the promoter was set to 1K upstream of TSS in *Arabidopsis*.

### Comparison analysis of wheat and *Arabidopsis*

The orthologous genes of wheat and *Arabidopsis* were downloaded from Ensembl BioMarts (Kinsella et al., 2011) (https://plants.ensembl.org/biomart/martview/5a1060ad127e25c7da355b64d87fe79a). DEGs in wheat and *Arabidopsis* were divided into three groups. Group I is species specific DEGs, that is, there are no orthologous genes in the other species. DEGs in Group II have orthologous genes in another species, but no expression or not DEGs. The orthologous genes of the Group III DEGs in another species are also DEGs.

Hierarchical clustering of wheat and *Arabidopsis* was performed as previously reported (Lemmon et al., 2016) in R. The TPM values of Group III DEGs and CPM values of ATAC-seq peaks located at the promoter and genic region of Group III DEGs were z-scaled within species, followed by hierarchical cluster analysis and principal component analysis.

To identify orthologous genes with similar and distinct expression patterns during regeneration of wheat and *Arabidopsis*, we performed pseudotime analysis in *Arabidopsis* following the method used in wheat. The Pearson correlation coefficient of gene expression between wheat and *Arabidopsis* was calculated and orthologous genes with R > 0.7 were thought to have similar expression patterns. Similar analysis were also performed with ATAC-seq data.

To compare TFs binding activity between wheat and *Arabidopsis* during regeneration, R package chromVAR (v1.10.0) (Schep et al., 2017) was used to calculate motif activity.

## Data availability

The raw sequence data reported in this paper have been deposited in the Genome Sequence Archive (Chen et al., 2021) in National Genomics Data Center (CNCB-NGDC Members and Partners, 2022), China National Center for Bioinformation / Beijing Institute of Genomics, Chinese Academy of Sciences (GSA: CRA008502) that are publicly accessible at https://ngdc.cncb.ac.cn/gsa.

## Acknowledgements

This research is supported by the Strategic Priority Research Program of the Chinese Academy of Sciences (XDA24010204) to J.X. and the National Natural Sciences Foundation of China (31730008) to X.S.Z., National Key Research and Development Program of China (2021YFD1201500) to J.X. and The Major Basic Research Program of Shandong Natural Science Foundation (ZR2019ZD15).

## Author contributions

J.X., X.-S. Z. and X.-M. L. designed and supervised the research and wrote the manuscript. X.-M. B. did the sample collection and in situ hybridization; M.-L. L. did plasmid construction and qRT-PCR. X.-M. B. and C.-Y. Z. did wheat transformation; X.-L. L. and X.-M. B. performed CUT&Tag, ATAC-seq and RNA-seq experiments; H.-Z, W. and X.-Y. Z did the reporter assay; X.-M. L. performed data analysis. X.-M. L., X.-M. B., Y.-M. Y and J.X. prepared all the figures. All authors discussed the results and commented on the manuscript.

## Extended Data Figure

**Extended Data Fig. 1.**
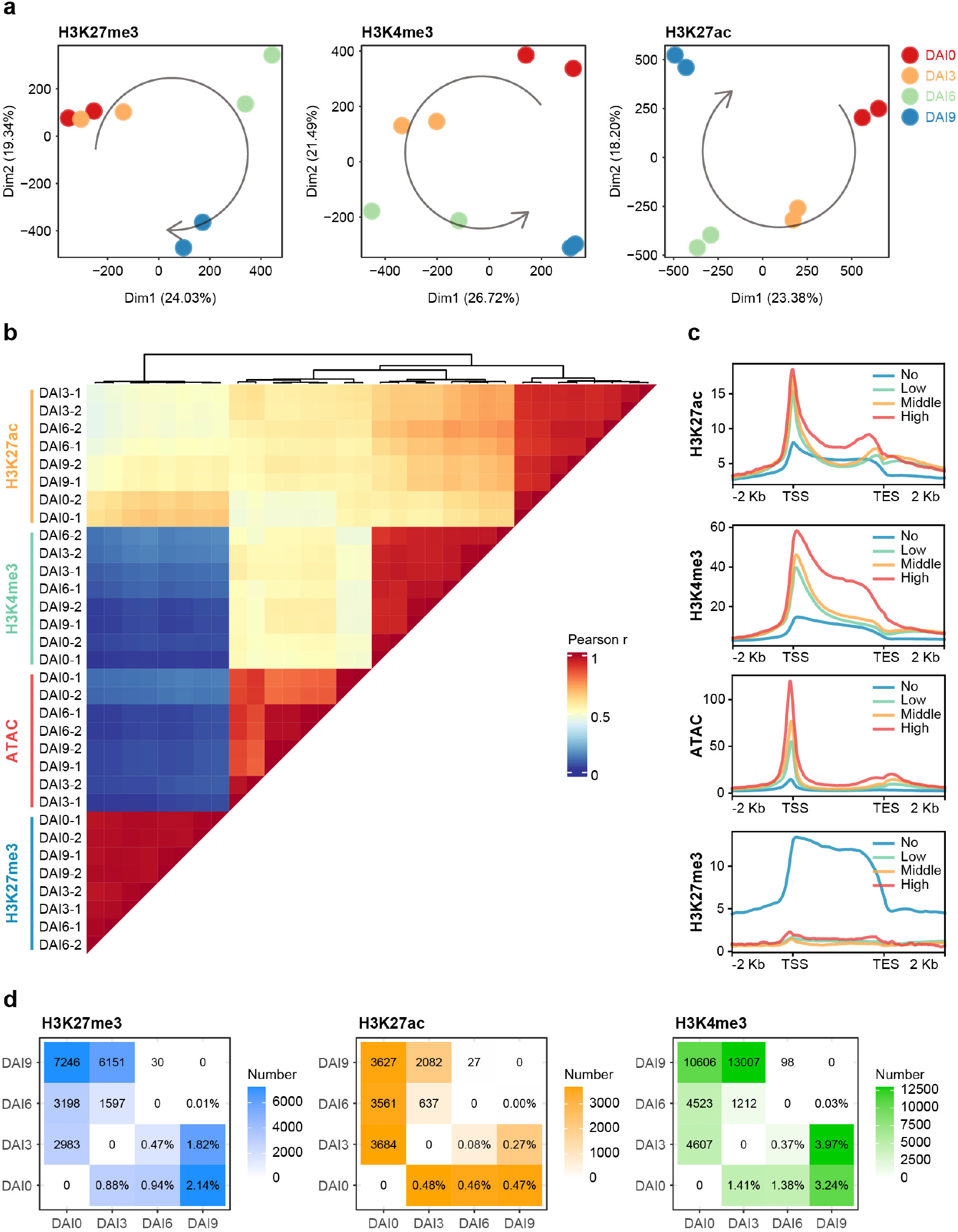
Overview of epigenome data of wheat shoot regeneration. **a**, Principal component analysis of H3K27me3, H3K4me3 and H3K27ac. **b**, Pearson correlation analysis of all epigenomic data. **c**, Epigenetic profiles on gene sets with different expression levels (data form DAI 6 stage); TSS: transcription start site, TES: transcription end site. **d**, Number of differentially marked peaks by H3K27me3, H3K27ac and H3K4me3 among different induction stages; Percentage represents the proportion of differentially histone modification peaks to all peaks.

**Extended Data Fig. 2.**
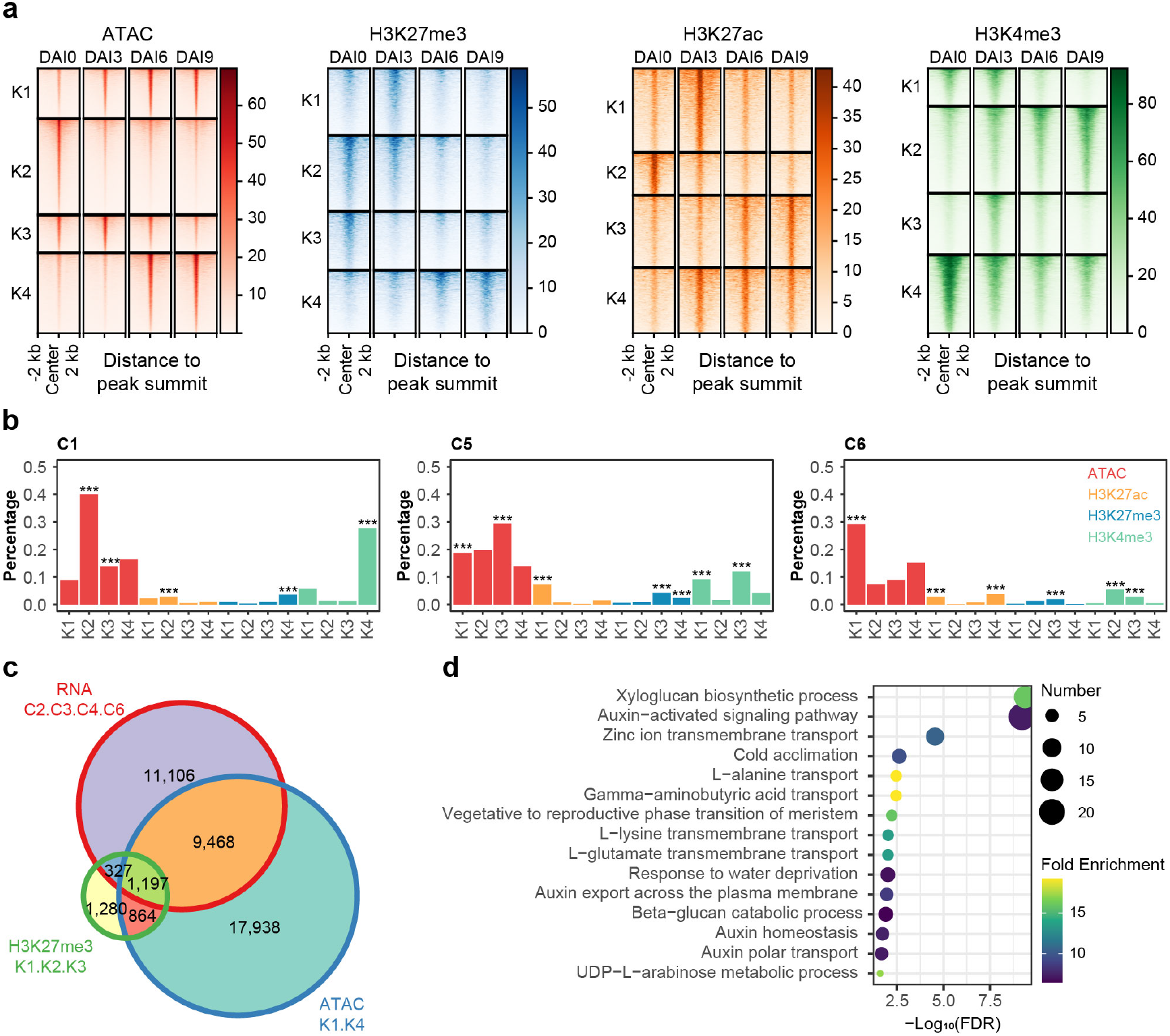
Chromatin dynamic during wheat shoot regeneration. **a**, K-means clustering analysis for differentially peaks of chromatin accessibility and histone modifications (H3K27me3, H3K27ac, H3K4me3). **b**, Over-representation analysis for gens in RNA cluster C1, C5 and C6 in ATAC and histone modification cluster (Fisher’s exact test, ***: p<0.001); The percentage represents the proportion of genes that overlap with the epigenetic cluster in the RNA cluster. **c**, Venn diagram for RNA cluster C2, C3, C4, C6, ATAC cluster K1, K4 and H3K27me3 cluster K1, K2, K3. **d**, The gene ontology enrichment analysis of 1,197 intersection genes in c.

**Extended Data Fig. 3.**
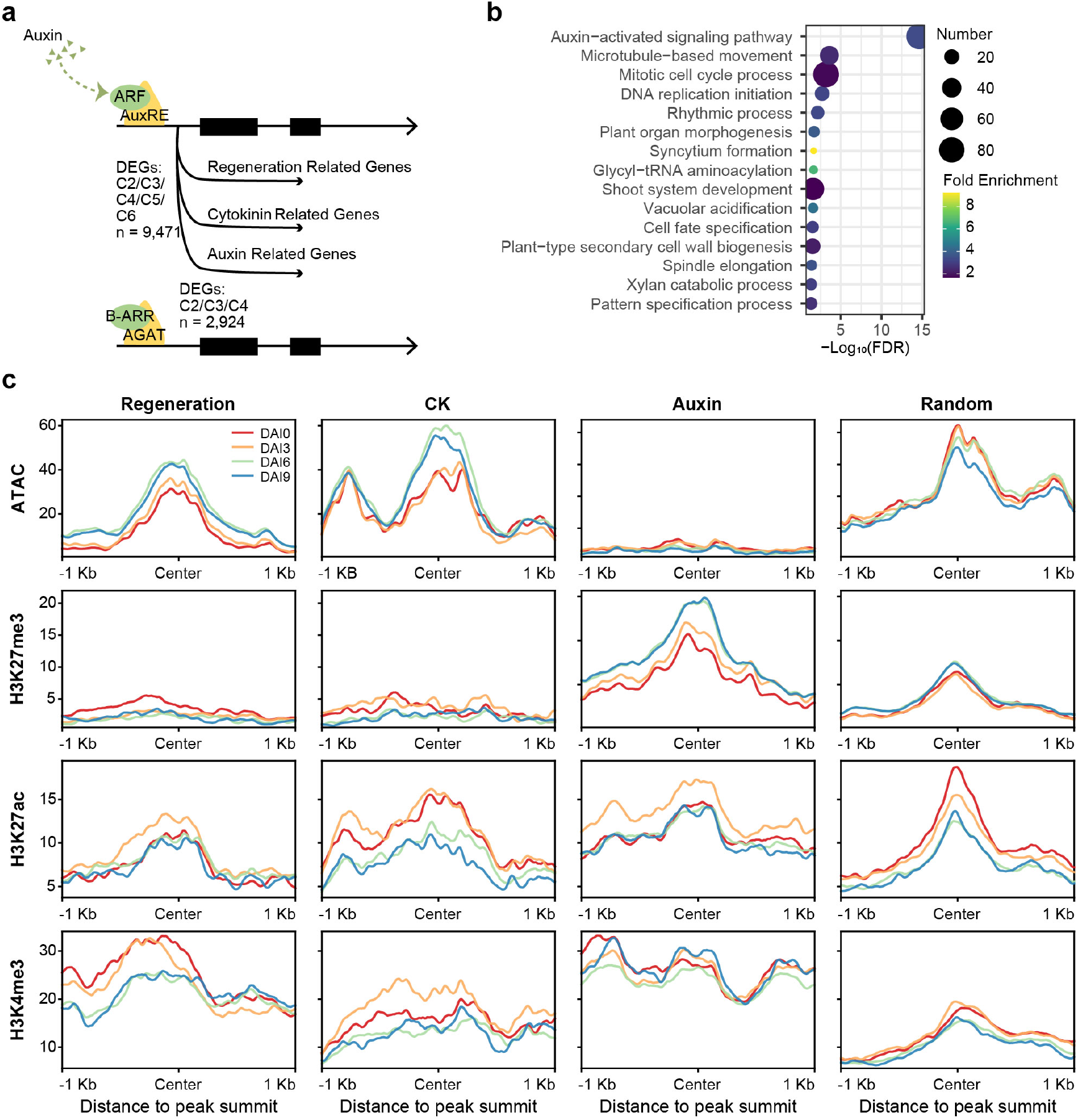
Comprehensive analysis of ARF target genes. **a**, Methods to identify ARF and type-B ARR target genes. **b**, Gene ontology enrichment analysis for ARFs target genes. **c**, Dynamic of chromatin accessibility and histone modification (H3K27me3, H3K27ac, H3K4me3) near AuxRE. The random background consists of randomly selected regions from other transcription factor binding sites.

**Extended Data Fig. 4.**
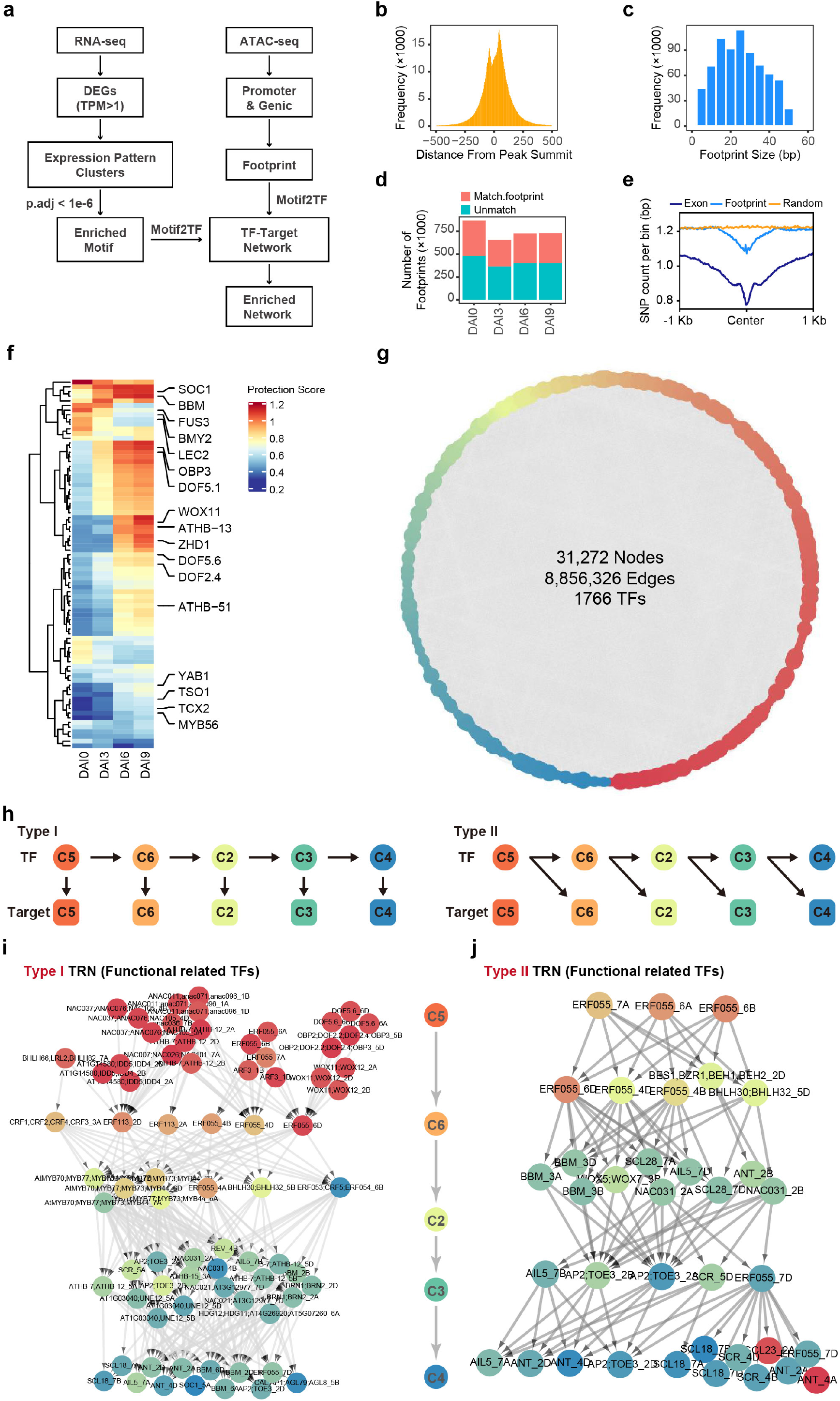
Construction of transcriptional regulatory network of wheat shoot regeneration. **a**, Pipeline for transcriptional regulatory network construction. **b**, Location distribution of footprint within the peak of ATAC-seq (data form DAI 6 stage). **c**, Distribution of the width of transcription factor footprint (data form DAI 6 stage). **d**, Numbers of footprint at different stages. **e**, Sequence conservation analysis at footprint sites; The background is randomly selected intergenic regions outside the open chromatin and coding regions; All regions are truncated to 25bp. **f**, Protection scores dynamic of differential footprint; Protection score reflects the chromatin accessibility of footprint. **g**, Overview of the transcriptional regulatory network of wheat shoot regeneration. **h**, Two transcriptional regulatory modes during wheat shoot regeneration. **i**, A network of functional related TFs in regulatory mode Type I. **j**, A network of functional related TFs in regulatory mode Type II.

**Extended Data Fig. 5.**
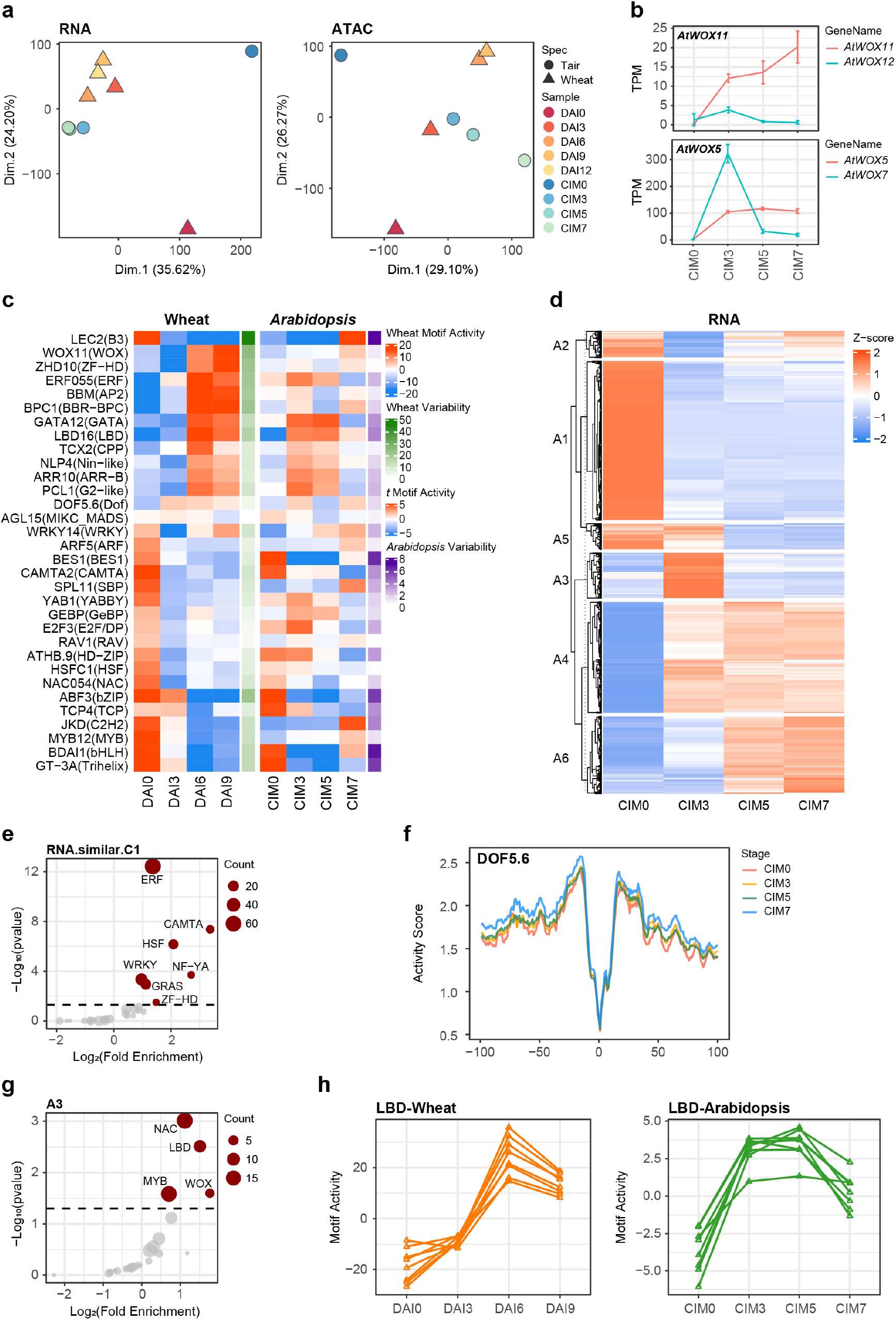
Comparative analysis of regeneration process between wheat and Arabidopsis. **a**, Principal component analysis of RNA-seq and ATAC-seq dataset during wheat and *Arabidopsis* regeneration. **b**, The expression pattern of *AtWOX11, AtWOX12, AtWOX5* and *AtWOX7* during *Arabidopsis* regeneration. **c**, Motif activity dynamic during regeneration in wheat and *Arabidopsis*; Motif activity represents the chromatin accessibility at the transcription factor binding sites. **d**, K-means clustering analysis of differentially expressed genes in *Arabidopsis*. **e**, TF family enrichment analysis of genes with similar expression pattern in C1. **f**, The footprint of DOF5.6 in *Arabidopsis* at different induction stages. **g**, TF family enrichment analysis of genes in *Arabidopsis* cluster A3. **h**, Motif activity dynamic of *LBDs* in wheat and *Arabidopsis*; Activity score reflects the chromatin accessibility.

**Extended Data Fig. 6.**
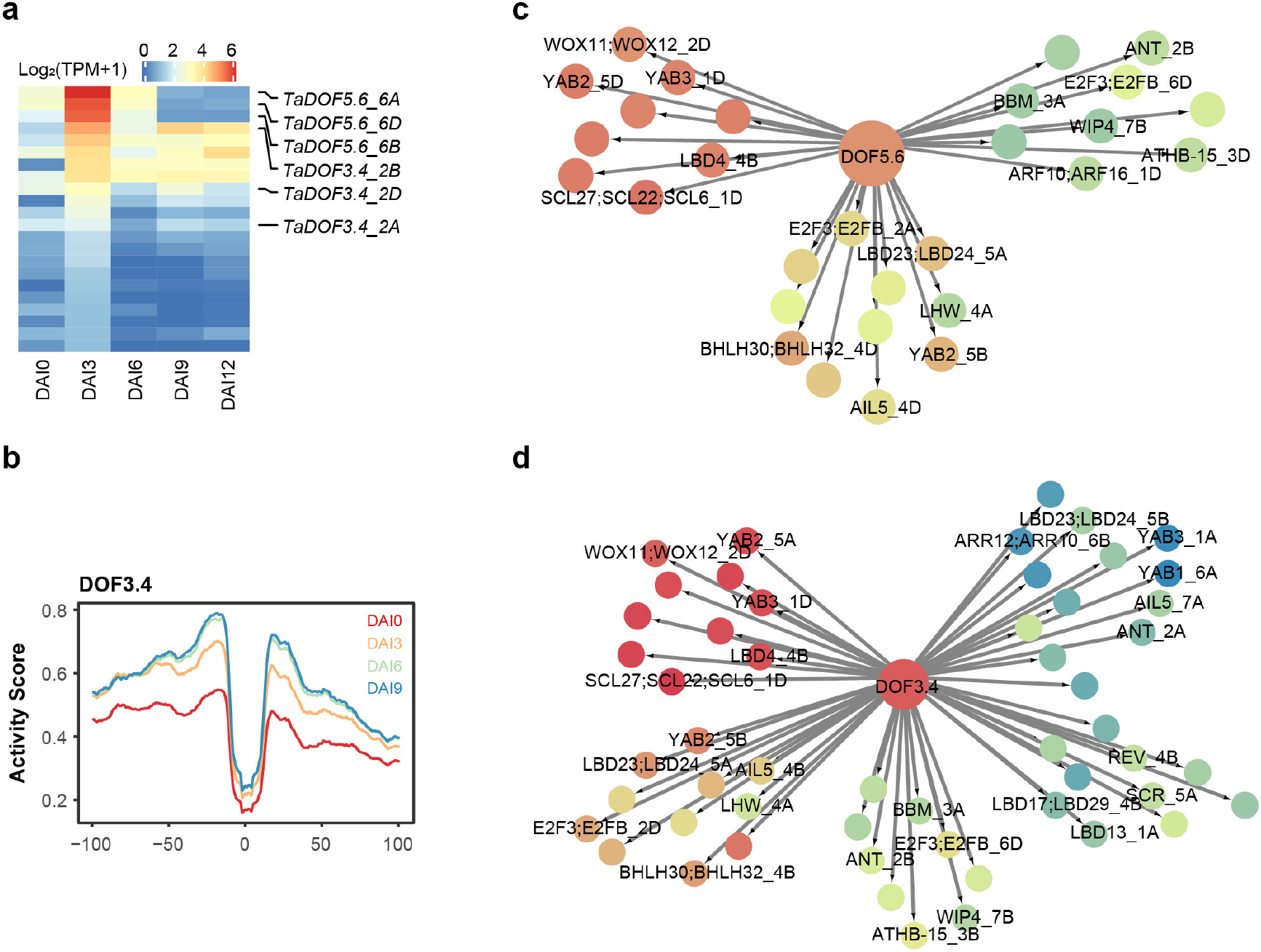
Expression and potential targets of DOFs during wheat shoot regeneration. **a**, The expression heatmap of DOF TFs during wheat shoot regeneration. The orthologous genes in *Arabidopsis* are shown. **b**, ATAC-seq footprint of DOF3.4 at different induction stages. **c**, The functional related target genes of *TaDOF5*.*6* in transcriptional regulatory network. **d**, The functional related target genes of *TaDOF3*.*4* in transcriptional regulatory network.

## References

Atta, R., Laurens, L., Boucheron-Dubuisson, E., Guivarc’h, A., Carnero, E., Giraudat-Pautot, V., Rech, P., and Chriqui, D. (2009). Pluripotency of Arabidopsis xylem pericycle underlies shoot regeneration from root and hypocotyl explants grown in vitro. The Plant Journal 57: 626–644.

Bidabadi, S.S. and Jain, S.M. (2020). Cellular, Molecular, and Physiological Aspects of In Vitro Plant Regeneration. Plants 9: 702.

Bie, X.M., Dong, L., Li, X.H., Wang, H., Gao, X.-Q., and Li, X.G. (2020). Trichostatin A and sodium butyrate promotes plant regeneration in common wheat. Plant Signaling & Behavior 15: 1820681.

Camacho, C., Coulouris, G., Avagyan, V., Ma, N., Papadopoulos, J., Bealer, K., and Madden, T.L. (2009). BLAST+: architecture and applications. BMC Bioinformatics 10: 421.

Chen, S., Zhou, Y., Chen, Y., and Gu, J. (2018). fastp: an ultra-fast all-in-one FASTQ preprocessor. Bioinformatics 34: i884–i890.

Chen, T. et al.. (2021). The Genome Sequence Archive Family: Toward Explosive Data Growth and Diverse Data Types. Genomics, Proteomics & Bioinformatics 19: 578–583.

CNCB-NGDC Members and Partners (2022). Database Resources of the National Genomics Data Center, China National Center for Bioinformation in 2022. Nucleic Acids Res 50: D27–D38.

Daimon, Y., Takabe, K., and Tasaka, M. (2003). The CUP-SHAPED COTYLEDON genes promote adventitious shoot formation on calli. Plant Cell Physiol 44: 113–121.

Danecek, P., Bonfield, J.K., Liddle, J., Marshall, J., Ohan, V., Pollard, M.O., Whitwham, A., Keane, T., McCarthy, S.A., Davies, R.M., and Li, H. (2021). Twelve years of SAMtools and BCFtools. GigaScience 10: giab008.

Debernardi, J.M., Tricoli, D.M., Ercoli, M.F., Hayta, S., Ronald, P., Palatnik, J.F., and Dubcovsky, J. (2020). A GRF-GIF chimeric protein improves the regeneration efficiency of transgenic plants. Nat Biotechnol 38: 1274–1279.

Della Rovere, F., Fattorini, L., D’Angeli, S., Veloccia, A., Falasca, G., and Altamura, M.M. (2013). Auxin and cytokinin control formation of the quiescent centre in the adventitious root apex of arabidopsis. Annals of Botany 112: 1395–1407.

Fan, M., Xu, C., Xu, K., and Hu, Y. (2012). LATERAL ORGAN BOUNDARIES DOMAIN transcription factors direct callus formation in Arabidopsis regeneration. Cell Res 22: 1169–1180.

Gordon, S.P., Heisler, M.G., Reddy, G.V., Ohno, C., Das, P., and Meyerowitz, E.M. (2007). Pattern formation during de novo assembly of the Arabidopsis shoot meristem. Development 134: 3539–3548.

Grant, C.E., Bailey, T.L., and Noble, W.S. (2011). FIMO: scanning for occurrences of a given motif. Bioinformatics 27: 1017–1018.

Gu, Z., Eils, R., and Schlesner, M. (2016). Complex heatmaps reveal patterns and correlations in multidimensional genomic data. Bioinformatics 32: 2847–2849.

H, L. and R, D. (2009). Fast and accurate short read alignment with Burrows-Wheeler transform. Bioinformatics (Oxford, England) 25.

Haberlandt, G. (2003). Culturversuche mit isolierten Pflanzenzellen. In Plant Tissue Culture: 100 years since Gottlieb Haberlandt, M. Laimer and W. Rücker, eds (Springer: Vienna), pp. 1–24.

Hao, C. et al.. (2020). Resequencing of 145 Landmark Cultivars Reveals Asymmetric Sub-genome Selection and Strong Founder Genotype Effects on Wheat Breeding in China. Molecular Plant 13: 1733–1751.

Hayta, S., Smedley, M.A., Demir, S.U., Blundell, R., Hinchliffe, A., Atkinson, N., and Harwood, W.A. (2019). An efficient and reproducible Agrobacterium-mediated transformation method for hexaploid wheat (Triticum aestivum L.). Plant Methods 15: 121.

He, C., Chen, X., Huang, H., and Xu, L. (2012). Reprogramming of H3K27me3 is critical for acquisition of pluripotency from cultured Arabidopsis tissues. PLoS Genet 8: e1002911.

Hiei, Y., Ishida, Y., and Komari, T. (2014). Progress of cereal transformation technology mediated by Agrobacterium tumefaciens. Frontiers in Plant Science 5.

Hoang, T. et al.. (2020). Gene regulatory networks controlling vertebrate retinal regeneration. Science 370: eabb8598.

Hu, B., Zhang, G., Liu, W., Shi, J., Wang, H., Qi, M., Li, J., Qin, P., Ruan, Y., Huang, H., Zhang, Y., and Xu, L. (2017). Divergent regeneration-competent cells adopt a common mechanism for callus initiation in angiosperms. Regeneration 4: 132–139.

Hu, X. and Xu, L. (2016). Transcription Factors WOX11/12 Directly Activate WOX5/7 to Promote Root Primordia Initiation and Organogenesis. Plant Physiol 172: 2363–2373.

Ikeuchi, M., Favero, D.S., Sakamoto, Y., Iwase, A., Coleman, D., Rymen, B., and Sugimoto, K. (2019). Molecular Mechanisms of Plant Regeneration. Annu Rev Plant Biol 70: 377–406.

International Wheat Genome Sequencing Consortium (IWGSC) (2018). Shifting the limits in wheat research and breeding using a fully annotated reference genome. Science 361: eaar7191.

Ishida, Y., Tsunashima, M., Hiei, Y., and Komari, T. (2015). Wheat (Triticum aestivum L.) transformation using immature embryos. Methods Mol Biol 1223: 189–198.

Jiang, F., Ryabova, D., Diedhiou, J., Hucl, P., Randhawa, H., Marillia, E.-F., Foroud, N.A., Eudes, F., and Kathiria, P. (2017). Trichostatin A increases embryo and green plant regeneration in wheat. Plant Cell Rep 36: 1701–1706.

Kareem, A., Durgaprasad, K., Sugimoto, K., Du, Y., Pulianmackal, A.J., Trivedi, Z.B., Abhayadev, P.V., Pinon, V., Meyerowitz, E.M., Scheres, B., and Prasad, K. (2015). PLETHORA Genes Control Regeneration by a Two-Step Mechanism. Current Biology 25: 1017–1030.

Kim, D., Paggi, J.M., Park, C., Bennett, C., and Salzberg, S.L. (2019). Graph-based genome alignment and genotyping with HISAT2 and HISAT-genotype. Nat Biotechnol 37: 907–915.

Kinsella, R.J., Kähäri, A., Haider, S., Zamora, J., Proctor, G., Spudich, G., Almeida-King, J., Staines, D., Derwent, P., Kerhornou, A., Kersey, P., and Flicek, P. (2011). Ensembl BioMarts: a hub for data retrieval across taxonomic space. Database (Oxford) 2011: bar030.

Klemm, S.L., Shipony, Z., and Greenleaf, W.J. (2019). Chromatin accessibility and the regulatory epigenome. Nat Rev Genet 20: 207–220.

Lee, K., Park, O.-S., Choi, C.Y., and Seo, P.J. (2019). ARABIDOPSIS TRITHORAX 4 Facilitates Shoot Identity Establishment during the Plant Regeneration Process. Plant Cell Physiol 60: 826–834.

Lee, K., Park, O.-S., Go, J.Y., Yu, J., Han, J.H., Kim, J., Bae, S., Jung, Y.J., and Seo, P.J. (2021). Arabidopsis ATXR2 represses de novo shoot organogenesis in the transition from callus to shoot formation. Cell Reports 37: 109980.

Lee, K., Park, O.-S., and Seo, P.J. (2017). Arabidopsis ATXR2 deposits H3K36me3 at the promoters of LBD genes to facilitate cellular dedifferentiation. Sci Signal 10: eaan0316.

Leiboff, S. and Hake, S. (2019). Reconstructing the Transcriptional Ontogeny of Maize and Sorghum Supports an Inverse Hourglass Model of Inflorescence Development. Current Biology 29: 3410-3419.e3.

Lemmon, Z.H., Park, S.J., Jiang, K., Van Eck, J., Schatz, M.C., and Lippman, Z.B. (2016). The evolution of inflorescence diversity in the nightshades and heterochrony during meristem maturation. Genome Res 26: 1676–1686.

Levin, M. et al.. (2016). The mid-developmental transition and the evolution of animal body plans. Nature 531: 637–641.

Li, Z., Schulz, M.H., Look, T., Begemann, M., Zenke, M., and Costa, I.G. (2019). Identification of transcription factor binding sites using ATAC-seq. Genome Biology 20: 45.

Liao, Y., Smyth, G.K., and Shi, W. (2014). featureCounts: an efficient general purpose program for assigning sequence reads to genomic features. Bioinformatics 30: 923–930.

Liu, H., Zhang, H., Dong, Y.X., Hao, Y.J., and Zhang, X.S. (2018a). DNA METHYLTRANSFERASE1-mediated shoot regeneration is regulated by cytokinininduced cell cycle in Arabidopsis. New Phytologist 217: 219–232.

Liu, J., Hu, X., Qin, P., Prasad, K., Hu, Y., and Xu, L. (2018b). The WOX11-LBD16 Pathway Promotes Pluripotency Acquisition in Callus Cells During De Novo Shoot Regeneration in Tissue Culture. Plant Cell Physiol 59: 734–743.

Liu, J., Sheng, L., Xu, Y., Li, J., Yang, Z., Huang, H., and Xu, L. (2014). WOX11 and 12 are involved in the first-step cell fate transition during de novo root organogenesis in Arabidopsis. Plant Cell 26: 1081–1093.

Liu, W. et al.. (2022). Transcriptional landscapes of de novo root regeneration from detached Arabidopsis leaves revealed by time-lapse and single-cell RNA sequencing analyses. Plant Communications 3: 100306.

Love, M.I., Huber, W., and Anders, S. (2014). Moderated estimation of fold change and dispersion for RNA-seq data with DESeq2. Genome Biology 15: 550.

Lowe, K. et al.. (2016). Morphogenic Regulators Baby boom and Wuschel Improve Monocot Transformation. Plant Cell 28: 1998–2015.

Meng, W.J., Cheng, Z.J., Sang, Y.L., Zhang, M.M., Rong, X.F., Wang, Z.W., Tang, Y.Y., and Zhang, X.S. (2017). Type-B ARABIDOPSIS RESPONSE REGULATORs Specify the Shoot Stem Cell Niche by Dual Regulation of WUSCHEL. Plant Cell 29: 1357–1372.

Mi, H., Muruganujan, A., Ebert, D., Huang, X., and Thomas, P.D. (2019). PANTHER version 14:pmore genomes, a new PANTHER GO-slim and improvements in enrichment analysis tools. Nucleic Acids Research 47: D419–D426.

Qiu, F., Xing, S., Xue, C., Liu, J., Chen, K., Chai, T., and Gao, C. (2022). Transient expression of a TaGRF4-TaGIF1 complex stimulates wheat regeneration and improves genome editing. Sci. China Life Sci. 65: 731–738.

Quinlan, A.R. and Hall, I.M. (2010). BEDTools: a flexible suite of utilities for comparing genomic features. Bioinformatics 26: 841–842.

Radhakrishnan, D., Kareem, A., Durgaprasad, K., Sreeraj, E., Sugimoto, K., and Prasad, K. (2018). Shoot regeneration: a journey from acquisition of competence to completion. Current Opinion in Plant Biology 41: 23–31.

Ramírez, F., Ryan, D.P., Grüning, B., Bhardwaj, V., Kilpert, F., Richter, A.S., Heyne, S., Dündar, F., and Manke, T. (2016). deepTools2: a next generation web server for deep-sequencing data analysis. Nucleic Acids Research 44: W160–W165.

Sarkar, A.K., Luijten, M., Miyashima, S., Lenhard, M., Hashimoto, T., Nakajima, K., Scheres, B., Heidstra, R., and Laux, T. (2007). Conserved factors regulate signalling in Arabidopsis thaliana shoot and root stem cell organizers. Nature 446: 811–814.

Schep, A.N., Wu, B., Buenrostro, J.D., and Greenleaf, W.J. (2017). chromVAR: inferring transcription-factor-associated accessibility from single-cell epigenomic data. Nat Methods 14: 975–978.

Schulze, S., Schäfer, B.N., Parizotto, E.A., Voinnet, O., and Theres, K. (2010). LOST MERISTEMS genes regulate cell differentiation of central zone descendants in Arabidopsis shoot meristems. Plant J 64: 668–678.

Shin, J., Bae, S., and Seo, P.J. (2019). De novo shoot organogenesis during plant regeneration. Journal of experimental botany 71.

Sugimoto, K., Jiao, Y., and Meyerowitz, E.M. (2010). Arabidopsis Regeneration from Multiple Tissues Occurs via a Root Development Pathway. Developmental Cell 18: 463–471.

Suo, J., Zhou, C., Zeng, Z., Li, X., Bian, H., Wang, J., Zhu, M., and Han, N. (2021). Identification of regulatory factors promoting embryogenic callus formation in barley through transcriptome analysis. BMC Plant Biol 21: 145.

Thorpe, T.A. (2007). History of plant tissue culture. Mol Biotechnol 37: 169–180.

Thorvaldsdóttir, H., Robinson, J.T., and Mesirov, J.P. (2013). Integrative Genomics Viewer (IGV): high-performance genomics data visualization and exploration. Briefings in Bioinformatics 14: 178–192.

Tian, F., Yang, D.-C., Meng, Y.-Q., Jin, J., and Gao, G. (2020). PlantRegMap: charting functional regulatory maps in plants. Nucleic Acids Research 48: D1104–D1113.

Wang, F.-X., Shang, G.-D., Wu, L.-Y., Xu, Z.-G., Zhao, X.-Y., and Wang, J.-W. (2020a). Chromatin Accessibility Dynamics and a Hierarchical Transcriptional Regulatory Network Structure for Plant Somatic Embryogenesis. Developmental Cell 54: 742-757.e8.

Wang, K., Liu, H., Du, L., and Ye, X. (2017). Generation of marker-free transgenic hexaploid wheat via an Agrobacterium-mediated co-transformation strategy in commercial Chinese wheat varieties. Plant Biotechnology Journal 15: 614–623.

Wang, K., Shi, L., Liang, X., Zhao, P., Wang, W., Liu, J., Chang, Y., Hiei, Y., Yanagihara, C., Du, L., Ishida, Y., and Ye, X. (2022). The gene TaWOX5 overcomes genotype dependency in wheat genetic transformation. Nat. Plants 8: 110–117.

Wang, W., Wang, Z., Li, X., Ni, Z., Hu, Z., Xin, M., Peng, H., Yao, Y., Sun, Q., and Guo, W. (2020b). SnpHub: an easy-to-set-up web server framework for exploring large-scale genomic variation data in the post-genomic era with applications in wheat. GigaScience 9.

Weijers, D. and Wagner, D. (2016). Transcriptional Responses to the Auxin Hormone. Annu Rev Plant Biol 67: 539–574.

Wu, L.-Y., Shang, G.-D., Wang, F.-X., Gao, J., Wan, M.-C., Xu, Z.-G., and Wang, J.-W. (2022). Dynamic chromatin state profiling reveals regulatory roles of auxin and cytokinin in shoot regeneration. Dev Cell 57: 526-542.e7.

Wu, T. et al.. (2021). clusterProfiler 4.0: A universal enrichment tool for interpreting omics data. Innovation 2.

Xu, M., Du, Q., Tian, C., Wang, Y., and Jiao, Y. (2021). Stochastic gene expression drives mesophyll protoplast regeneration. Science Advances 7: eabg8466.

Yu, G., Wang, L.-G., and He, Q.-Y. (2015). ChIPseeker: an R/Bioconductor package for ChIP peak annotation, comparison and visualization. Bioinformatics 31: 2382–2383.

Zhang, W., Yin, M.-Q., Zhao, P., Wang, K., Du, L.-P., and Ye, X.-G. (2018). Regeneration Capacity Evaluation of Some Largely Popularized Wheat Varieties in China. Acta Agronomica Sinica 44: 208.

Zhang, Y., Liu, T., Meyer, C.A., Eeckhoute, J., Johnson, D.S., Bernstein, B.E., Nusbaum, C., Myers, R.M., Brown, M., Li, W., and Liu, X.S. (2008). Model-based analysis of ChIP-Seq (MACS). Genome Biol 9: R137.

Zhao, L., Lin, X., Yang, Y., Bie, X., Zhang, H., Chen, J., Liu, X., Wang, H., Jiang, J., Fu, X., Zhang, X., and Xiao, J. (2022). Chromatin reprogramming and transcriptional regulation orchestrate embryogenesis in hexaploid wheat.: 2022.01.21.477188.

Zhao, N., Zhang, K., Wang, C., Yan, H., Liu, Y., Xu, W., and Su, Z. (2020). Systematic Analysis of Differential H3K27me3 and H3K4me3 Deposition in Callus and Seedling Reveals the Epigenetic Regulatory Mechanisms Involved in Callus Formation in Rice. Frontiers in Genetics 11.

Zhou, Y. et al.. (2020). Triticum population sequencing provides insights into wheat adaptation. Nat Genet 52: 1412–1422.

